# Directed Evolution of a Genetically Encoded Indicator for Chloride

**DOI:** 10.1101/2024.11.25.624492

**Authors:** Weicheng Peng, Jasmine N. Tutol, Shelby M. Phelps, Hiu Kam, Jacob K. Lynd, Sheel C. Dodani

**Affiliations:** Departments of Chemistry and Biochemistry, The University of Texas at Dallas, Richardson, TX 75080; Biological Sciences, The University of Texas at Dallas, Richardson, TX 75080

**Keywords:** chloride, fluorescent indicator, GFP, protein engineering

## Abstract

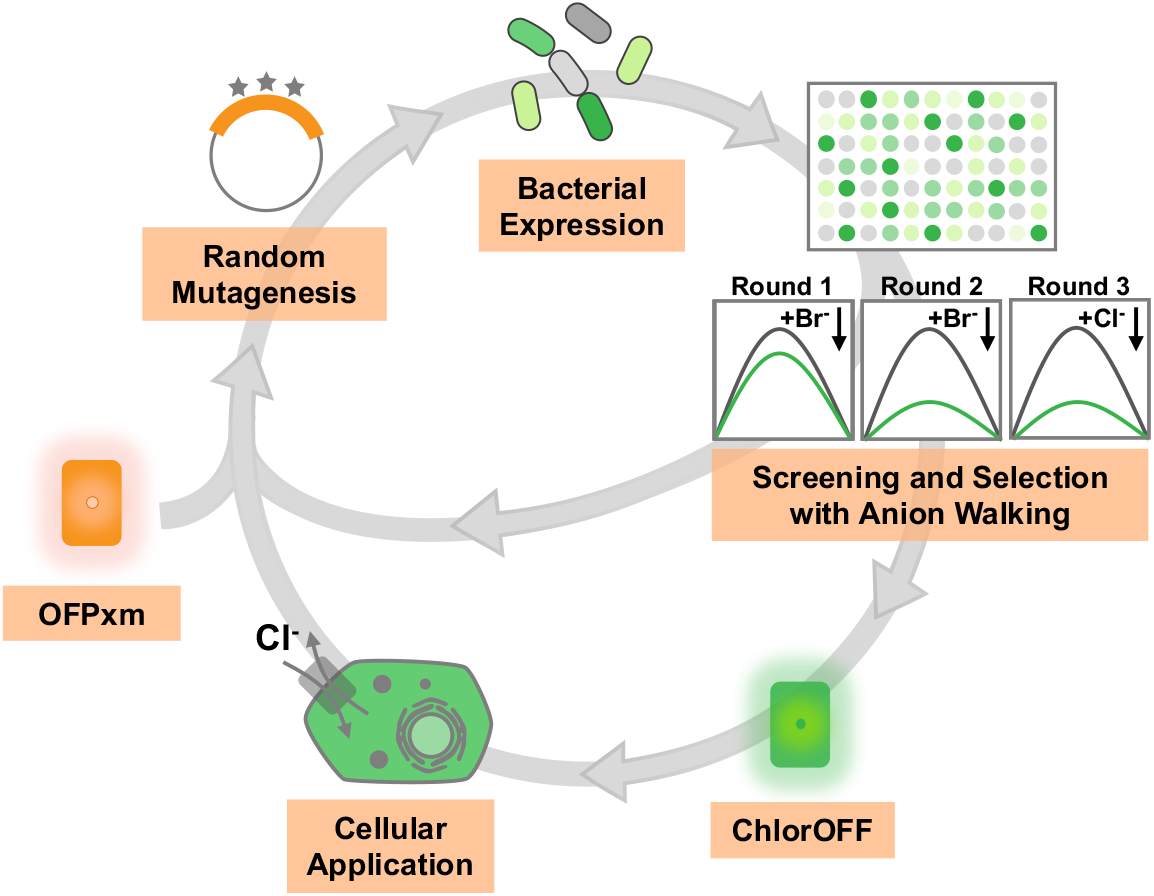

Inarguably, the green fluorescent protein (GFP) family is an exemplary model for protein engineering, accessing a range of unparalleled functions and utility in biology. The first variant to recognize and provide an optical output of chloride in living cells was serendipitously uncovered more than 25 years ago. Since then, researchers have actively expanded the potential of GFP indicators for chloride through site-directed and combinatorial site-saturation mutagenesis, along with chimeragenesis. However, to date, the power of directed evolution has yet to be unleashed. As a proof-of-concept, here, we use random mutagenesis paired with anion walking to engineer a chloride-insensitive fluorescent protein named OFPxm into a functional indicator named ChlorOFF. The sampled mutational landscape unveils an evolutionary convergent solution at one position in the anion binding pocket and nine other mutations across eight positions, of which only one has been previously linked to chloride sensing potential in the GFP family.

## Main Text

The green fluorescent protein (GFP) was unearthed from jellyfish *Aequorea victoria* more than 50 years ago and is still considered a “guiding star” biotechnology.^1,2^ Historically, GFP has proven to be mutationally resilient, unveiling new mechanistic insights and functionality potential.^2^ Structurally, the β-barrel encases an intrinsic chromophore on an internal α-helix (Figure S1).^3,4^ The chromophore results from the autocatalytic cyclization, followed by dehydration and oxidation of three amino acids (S65/Y66/G67). A break in the hydrogen bonding pattern occurs near the chromophore in a region defined by the gate post (Y145/H148) and β-bulge (N146/S147) on the β7 strand. The chromophore is exposed to bulk water through a pore between the β7 and β10 strands.

In the context of GFP, mutations within and around the chromophore not only shift the spectral profile but also serendipitously unlock sensitivity to anions such as chloride.^5-7^ One pioneering variant is known as avYFP-H148Q (Figure 1A; Figure S2).^5–7^. Since chloride is the most abundant biological anion, researchers have actively sought to engineer this function for cellular imaging applications.^8^ To this end, site-directed and combinatorial site-saturation mutagenesis, along with chimeragenesis to other fluorescent proteins, have expanded the chloride sensing potential of avYFP-H148Q with further extensions to the GFP variants E^2^GFP and Topaz.^9–19^

**Figure 1.**
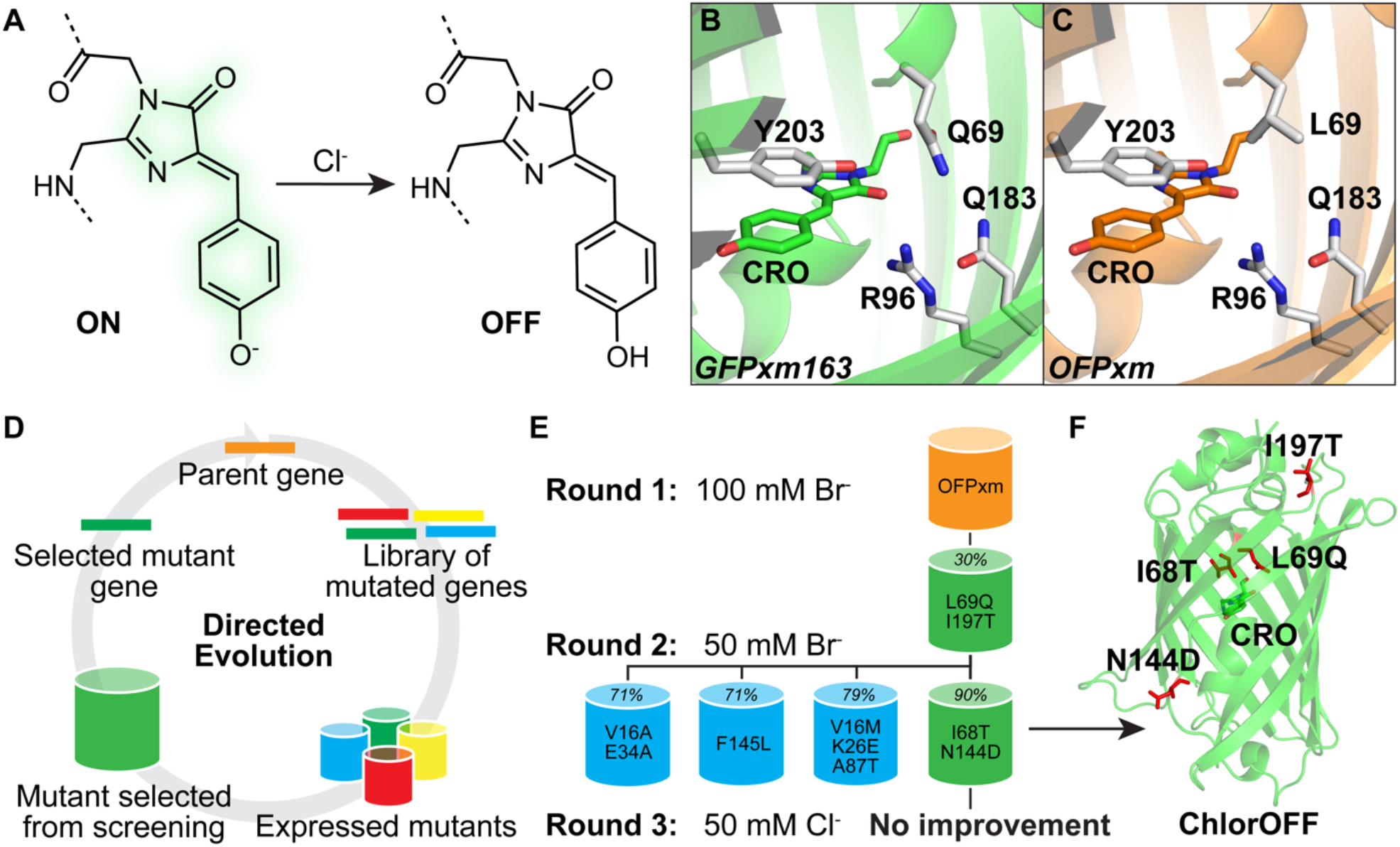
Engineering of ChlorOFF from the chloride-insensitive fluorescent protein OFPxm. (A) Chloride (Cl^-^) sensing mechanism for avYFP-like indicators. Homology models of (B) GFPxm163 and (C) OFPxm with the putative anion binding pocket near the chromophore (CRO). (D) General directed evolution workflow. (E) Selected mutants from each round of directed evolution are listed with the turn-off response with bromide (Br^-^) from the lysate rescreen (Figure S10). (F) Homology model of ChlorOFF with all mutated residues shown as red sticks.

Beyond avGFP, new templates have been sourced from the GFP family. One such example is the red fluorescent protein mBeRFP derived from EqFP578 in *Entacmaea quadricolor*.^20^ Rational transplantation of mutations from avYFP-H148Q variants and E^2^GFP enhanced chloride sensing. As a complementary approach, we have relied on structure-guided bioinformatics, drawing focus to the chloride coordination sphere, to uncover chloride-sensitive members. Both phiYFP in *Phialidium sp*. SL-2003 and GFPxm163 from GFPxm in *Aequorea macrodactyla* have avYFP-like binding pockets, whereas mNeonGreen from LanYFP in the *Branchiostoma lanceolatum* is unique.^21–23^ Of these, mNeonGreen has been further engineered into the ChlorONs using combinatorial site-saturation mutagenesis in the binding pocket.^24^

Looking to the progress over the last 25 years, the power of directed evolution has yet to be applied to engineer fluorescent protein indicators for chloride.^25^ In this Letter, we have explored if the directed evolution of a chloride-insensitive fluorescent protein with random mutagenesis could give rise to a chloride-sensitive variant with comparable or enhanced function to the current state of the art, and even unveil mutational hotspots. As a proof-of-concept, a variant of GFPxm named OFPxm was selected.^26^ Relative to GFPxm163 described above, OFPxm is 98% identical as it bears only four mutations (Figure 1B, 1C; Figure S3). Of these, three are on the N-terminal β1 strand (I11V, V14I, I16V), and one is in the avYFP-like binding pocket (Q69L). Notably, some mutations at position 69 in avYFP-H148Q and phiYFP do not affect chromophore maturation, despite proximity, but can lower or eliminate sensitivity to anions.^10,27^

With this logic, we first tested if OFPxm would be insensitive to chloride. To do so, it was expressed, purified, and spectroscopically evaluated (Figure S4–S7). Based on the absorption spectra, the tyrosine-based chromophore undergoes a pH-dependent equilibrium between the phenol (*λ*_abs_ = 400 nm) and phenolate (*λ*_abs_ = 512 nm with a shoulder at *λ*_abs_ = 486 nm) states (p*K*_a_ = 6.49 ± 0.03). While the phenol state is non-emissive, the phenolate state has an emission maximum centered at 524 nm. As expected, no spectral changes are observed upon the addition of 100 mM chloride (p*K*_a_ = 6.50 ± 0.02). Thus, confirming the amino acid differences between OFPxm and GFPxm163 could contribute to the chloride sensing phenotype.

With this starting point, we developed a directed evolution workflow paired with an anion walking screening and selection strategy to introduce chloride sensitivity into OFPxm (Figure 1D). The error-prone polymerase chain reaction (PCR) was used to introduce random mutations in the coding sequence (Table S1–S3).^25^ Following initial assay optimization, approximately (*ca*.) 1,800 *E. coli* colonies were expressed in a 96-well format for each round (Figure S8, S9). The fluorescence response of each variant was screened in cell lysate in the absence and presence of anion at pH 7. Initially, bromide was used as a surrogate for chloride because it has a larger ionic radius (Br^-^ 2 Å > Cl^-^ 1.8 Å) and lower dehydration enthalpy (Br^-^ 335 kJ/mol < Cl^-^ 365 kJ/mol).^28^ These physical differences can translate to a higher affinity complex for bromide, resulting in a greater fluorescence response.^24^ Based on this rationale, we used 100 mM bromide in the first round and selected the L69Q/I197T variant with 30% fluorescence quenching (Figure 1E; Figure S10, S11). With this initial gain, the selection pressure was increased to 50 mM bromide in the second round to enhance sensitivity. This resulted in the identification of four variants bearing the following additional mutations: I68T/N144D (90 ± 2%) > V16M/K26E/A87T (79 ± 3 %) > F145L (71 ± 4%) = V16A/E34V (71 ± 2%) (Figure 1E; Figure S10, S11). Since the I68T/L69Q/N144D/I197T variant had the largest degree of fluorescence quenching with bromide, it was selected as the parent for the third round with a shift to 50 mM chloride for the screening assay. However, no improvements were observed, yielding the final variant – OFPxm-I68T/L69Q/N144D/I197T – that we named ChlorOFF and selected for further investigation in purified form (Figure 1E, 1F; Figure S10–S13).

Unsurprisingly, apo ChlorOFF has a pH-dependent absorption profile between the phenol (*λ*_abs_ = 400 nm) and phenolate (*λ*_abs_ = 512 nm with a shoulder at 486 nm) chromophore states (Figure S14, S15). However, the addition of 100 mM chloride increases the chromophore p*K*_a_ from 6.37 ± 0.04 to 8.00 ± 0.01, favoring the phenol state. In line with these data, chloride can tune the chromophore equilibrium in a dose-dependent fashion under constant pH conditions (Figure 2A; Figure S16–S18). Specifically at pH 7, apo ChlorOFF has a bright phenolate-centered emission maximum (*λ*_em_ = 524 nm, ε×QY = 6.19 ± 0.29) that does not shift but is quenched by 90.3 ± 0.3% (ε×QY = could not be determined) with up to 128 mM chloride but not with gluconate as an ionic strength control (Figure 2B, 2C; Figure S19–S22; Table S4). Further fitting of these data to a single-site binding model indicates that the chloride binding affinity (*K*_d_) for ChlorOFF is 12.7 ± 0.2 mM (Figure S16; Table S4). Indeed, the overall brightness and emission response of ChlorOFF is pH-dependent but maintained across physiological regime from pH 6.5 (ε×QY = 4.96 ± 0.06, *K*_d_ = 6.03 ± 0.10 mM) to pH 7.5 (ε×QY = 7.91 ± 0.42, *K*_d_ = 31.6 ± 0.7 mM) (Figure S17–S21; Table S4). Finally, given our anion walking screening and selection strategy, we tested if ChlorOFF could also be sensitive to bromide, iodide, and nitrate at pH 7. Like chloride, all three anions can tune the chromophore equilibrium, resulting in comparable fluorescence quenching (> 95%) with relatively higher binding affinities: iodide (0.43 ± 0.02 mM) > nitrate (1.83 ± 0.05 mM) > bromide (4.63 ± 0.12 mM) (Figure 2C; Figure S23–25; Table S4).

**Figure 2.**
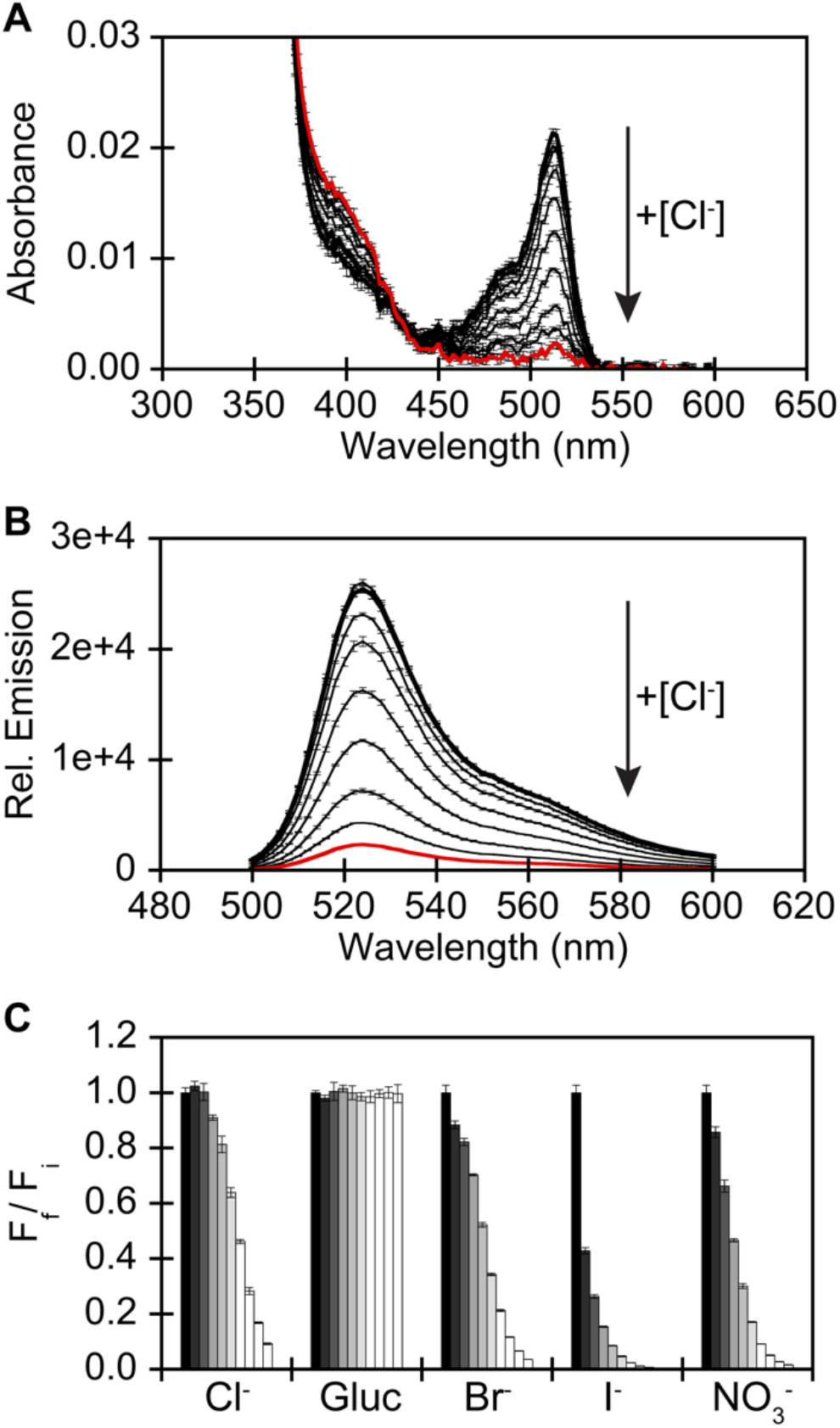
*In vitro* spectroscopic characterization of ChlorOFF. (A) Absorbance (*λ*_abs_ = 350–600 nm) and (B) emission (*λ*_ex_ = 480 nm, *λ*_em_ = 500–600 nm) spectra of ChlorOFF titrated with 0 mM (bold black) to 128 mM (red) chloride (Cl^-^). (C) Fluorescence response (*λ*_em_ = 524 nm) of ChlorOFF titrated with 0 mM (black) to 128 mM (white) chloride, gluconate (Gluc), bromide (Br^-^), iodide (I^-^), and nitrate (NO_3-_). All experiments were carried out at room temperature (24–26 °C) with 3 µM protein in 50 mM sodium phosphate buffer at pH 7. Data is shown for one protein batch (n = 3, average with standard error of the mean). The data for the second protein batch is shown in Figure S16.

Building from the *in vitro* spectroscopy, we evaluated the performance of ChlorOFF with fluorescence microscopy in the U-2 OS cell line, which is widely used for imaging applications given its epithelial morphology and ease for imaging.^29,30^ More importantly, for our goal, these osteosarcoma cells also express endogenous chloride-transporting proteins, thus bypassing the need for exogenous overexpression.^31,32^ For all experiments, a stable cell line expressing ChlorOFF in the cytosol and nucleus was generated over two cycles of enrichment (Figure S26). Cells imaged in the chloride-containing buffer have a detectable fluorescence signal (Figure S27, Movie S1). By relying on the inherent promiscuity of the indicator and chloride-transporting proteins, the extracellular chloride (137 mM) was partially exchanged with iodide (∼69 mM), resulting in further quenching (*ca*. 78% ± 4) (Figure S28, Movie S1). Given this result, we were motivated to probe if ChlorOFF could provide a direct readout of chloride transport, bypassing the need for an iodide surrogate. Looking to electrophysiology and conductivity assays, the bioavailable chloride pool was depleted by acute incubation in gluconate-containing buffer (137 mM). The fluorescence signal quenches (*ca*. 38% ± 13) upon replenishment of extracellular chloride (137 mM), but not with gluconate (*ca*. 1.17-fold ± 0.1) (Figure 3; Movie S2, S3). For these time-lapse exchange assays, the intracellular pH was monitored with BCECF with minimal detectable changes (Figure S29–S32, Movies S4–S6).^33^

**Figure 3.**
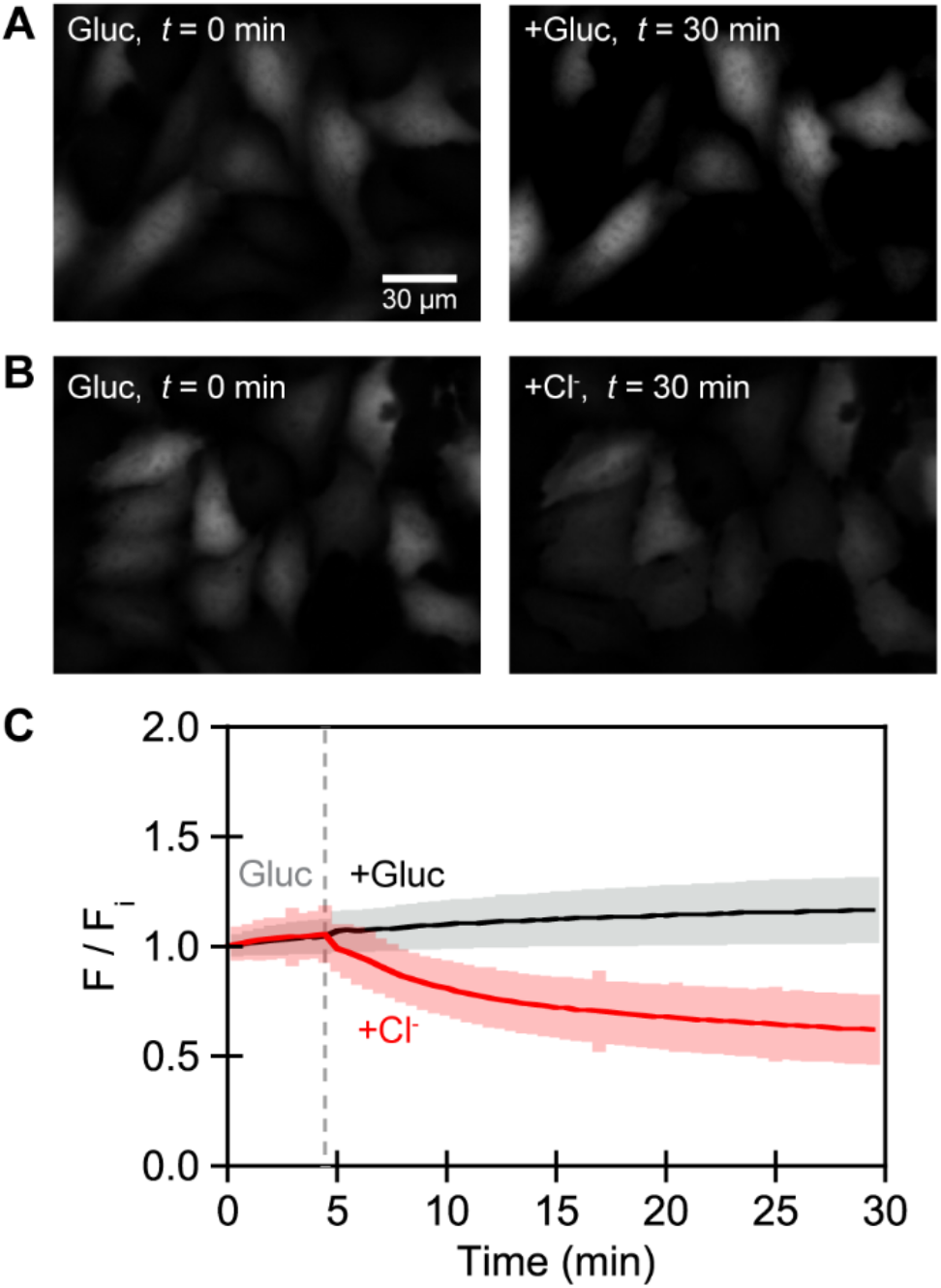
Characterization of ChlorOFF in the U-2 OS cell line with widefield fluorescence microscopy. Representative fluorescence images for the exchange assay with (a) gluconate-gluconate (*n* = 2,587) and (b) gluconate-chloride (*n* = 2,720). (c) For each condition, the average median fluorescence intensity with standard deviation (shaded) is reported for the *n* regions of interest from three biological replicates. The grey dashed line is the start of the perfusion for the anion exchange. All experiments were conducted at 37 °C in a PBS buffer at pH 7.4 (see *Methods*) containing 137 mM NaCl or NaGluc (Movie S2, S3). Abbreviation: Gluc, gluconate.

In summary, we have successfully demonstrated how directed evolution can be an effective strategy to unlock chloride sensitivity in a fluorescent protein. This rare demonstration with OFPxm resulted in the functional indicator ChlorOFF. Across the mutational landscape from OFPxm to ChlorOFF, L69Q provides a critical coordinating residue. Given the context of nearby residues, this mutation is rather unsurprising. Its essentiality is highlighted by evolutionary convergence of Q69 found in avYFP-H148Q, phiYFP, and GFPxm163. Next to L69Q, the chromophore is covalently anchored through I68T, which is linked to efficient protein folding, chromophore p*K*_a_, and chloride binding affinity.^7,34^ Aside from positions 68 and 69, the other sampled positions have yet to be connected to chloride sensing potential but could also have secondary effects (Table S5, Figure S33–37). In proximity to the chromophore, N144D and neighboring F145L were independently identified on a flexible loop that bears the gate post (positions 145 and 148) and β-bulge (positions 146 and 147) regions.^35^ Indeed, mutations along this loop can directly alter accessibility to water and even anions, thereby tuning photophysical properties.^7,36,37^ Except for V16M/A, the remaining mutations – K26E, E34V, A87T, and I197T – are solvent-exposed, positioned distally away from the chromophore and binding pocket. While we cannot precisely trace the functional significance of these positions to anion sensitivity, they could be explored as mutational hotspots down the line. Given that ChlorOFF retains an avYFP-like binding pocket, it has a similar p*K*_a_-dependent sensing mechanism and anion preference.^7^ However, the mutations in ChlorOFF increase the affinity for chloride by 18-fold relative to GFPxm163 – its nearest anion-sensitive neighbor – that does not bear any of the substitutions aside from Q69 (Figure S3).^22^ This engineered property translates into a direct readout of not only iodide, as expected, but also chloride transport in living cells. Building from this proof-of-concept, future endeavors will capitalize on the power of directed evolution – from lysate, in-cell, and *in silico* – to unleash the untapped potential of GFP indicators for chloride.^38,39^

## Methods

All experimental methods and data analysis are described in the Supporting Information.

## Supporting information

Fluorescence imaging movies

## Data Availability Statement

All the data in this study is available in the Main Text and Supporting Information. The corresponding author can be contacted for additional requests.

## Supporting Information

Experimental methods and data (PDF). Fluorescence imaging movies (AVI).

## Author contributions

S.C.D. designed and supervised the research project. W.P. carried out all *in vitro* experiments with analysis and preliminary in-cell imaging experiments. J.N.T. and S.M.P. carried out all in-cell imaging experiments with analysis. H.K. and J.K.L. carried out preliminary spectroscopy experiments. W.P., J.N.T., S.M.P., and S.C.D. wrote the manuscript with input from all the authors.

## Acknowledgements

J.N.T. acknowledges the Irving S. Sigal Postdoctoral Fellowship from the American Chemical Society. S.C.D. acknowledges the Welch Foundation (AT-1918-20170325, AT-206020210327) and the National Institute of General Medical Sciences of the National Institutes of Health (R35GM128923). This study does not represent the views of the funding agencies and is the sole responsibility of the authors.

## Conflicts of Interest

There are no conflicts to declare.

## Notes

### Competing Interest Statement

The authors have declared no competing interest.

## References

(1) The Nobel Prize in Chemistry 2008. NobelPrize.org.

(2) Tsien, R. Y. (1998) The green fluorescent protein. Annu. Rev. Biochem. 67, 509–544.

(3) Ormö, M., Cubitt, A. B., Kallio, K., Gross, L. A., Tsien, R. Y., and Remington, S. J. (1996) Crystal structure of the Aequorea victoria green fluorescent protein. Science 273, 1392–1395.

(4) Yang, F., Moss, L. G., and Phillips, G. N. (1996) The molecular structure of green fluorescent protein. Nat. Biotechnol. 14, 1246–1251.

(5) Wachter, R. M., and Remington, S. J. (1999) Sensitivity of the yellow variant of green fluorescent protein to halides and nitrate. Curr. Biol. 9, R628–R629.

(6) Jayaraman, S., Haggie, P., Wachter, R. M., Remington, S. J., and Verkman, A. S. (2000) Mechanism and cellular applications of a green fluorescent protein-based halide sensor. J. Biol. Chem. 275, 6047–6050.

(7) Wachter, R. M., Yarbrough, D., Kallio, K., and Remington, S. J. (2000) Crystallographic and energetic analysis of binding of selected anions to the yellow variants of green fluorescent protein. J. Mol. Biol. 301, 157–171.

(8) Lodovichi, C., Ratto, G. M., Trevelyan, A. J., and Arosio, D. (2022) Genetically encoded sensors for chloride concentration. J. Neurosci. Methods. 368, 109455.

(9) Shariati, K., Zhang, Y., Giubbolini, S., Parra, R., Liang, S., Edwards, A., Hejtmancik, J. F., Ratto, G. M., Arosio, D., and Ku, G. (2022) A superfolder green fluorescent protein-based biosensor allows monitoring of chloride in the endoplasmic reticulum. ACS Sens. 7, 2218–2224.

(10) Grimley, J. S., Li, L., Wang, W., Wen, L., Beese, L. S., Hellinga, H. W., and Augustine, G. J. (2013) Visualization of synaptic inhibition with an optogenetic sensor developed by cell-free protein engineering automation. J. Neurosci. 33, 16297–16309.

(11) Kuner, T., and Augustine, G. J. (2000) A genetically encoded ratiometric indicator for chloride: capturing chloride transients in cultured hippocampal neurons. Neuron. 27, 447–459.

(12) Markova, O., Mukhtarov, M., Real, E., Jacob, Y., and Bregestovski, P. (2008) Genetically encoded chloride indicator with improved sensitivity. J. Neurosci. Methods. 170, 67–76.

(13) Zhong, S., Navaratnam, D., and Santos-Sacchi, J. (2014) A genetically-encoded YFP sensor with enhanced chloride sensitivity, photostability and reduced pH interference demonstrates augmented transmembrane chloride movement by gerbil prestin (SLC26a5). PLOS ONE. 9, e99095.

(14) Galietta, L. J. V., Haggie, P. M., and Verkman, A. S. (2001) Green fluorescent protein-based halide indicators with improved chloride and iodide affinities. FEBS letters. 499, 220–224.

(15) Arosio, D., Garau, G., Ricci, F., Marchetti, L., Bizzarri, R., Nifosì, R., and Beltram, F. (2007) Spectroscopic and structural study of proton and halide ion cooperative binding to gfp. Biophys. J. 93, 232–244.

(16) Mukhtarov, M., Liguori, L., Waseem, T., Rocca, F., Buldakova, S., Arosio, D., and Bregestovski, P. (2013) Calibration and functional analysis of three genetically encoded Cl(^-^)/pH sensors. Front. Mol. Neurosci. 6, 9.

(17) Paredes, J. M., Idilli, A. I., Mariotti, L., Losi, G., Arslanbaeva, L. R., Sato, S. S., Artoni, P., Szczurkowska, J., Cancedda, L., Ratto, G. M., Carmignoto, G., and Arosio, D. (2016) Synchronous bioimaging of intracellular pH and chloride based on LSS fluorescent protein. ACS Chem. Biol. 11, 1652–1660.

(18) Arosio, D., Ricci, F., Marchetti, L., Gualdani, R., Albertazzi, L., and Beltram, F. (2010) Simultaneous intracellular chloride and pH measurements using a GFP-based sensor. Nat. Methods. 7, 516–518.

(19) Raimondo, J. V., Joyce, B., Kay, L., Schlagheck, T., Newey, S. E., Srinivas, S., and Akerman, C. J. (2013) A genetically-encoded chloride and pH sensor for dissociating ion dynamics in the nervous system. Front. Cell. Neurosci. 7, 202.

(20) Salto, R., Giron, M. D., Puente-Muñoz, V., Vilchez, J. D., Espinar-Barranco, L., Valverde-Pozo, J., Arosio, D., and Paredes, J. M. (2021) New red-emitting chloride-sensitive fluorescent protein with biological uses. ACS Sens. 6, 2563–2573.

(21) Tutol, J. N., Peng, W., and Dodani, S. C. (2019) Discovery and characterization of a naturally occurring, turn-on yellow fluorescent protein sensor for chloride. Biochemistry. 58, 31–35.

(22) Peng, W., Maydew, C. C., Kam, H., Lynd, J. K., Tutol, J. N., Phelps, S. M., Abeyrathna, S., Meloni, G., and Dodani, S. C. (2022) Discovery of a monomeric green fluorescent protein sensor for chloride by structure-guided bioinformatics. Chem. Sci. 13, 12659–12672.

(23) Tutol, J. N., Kam, H. C., and Dodani, S. C. (2019) Identification of mNeonGreen as a pH-dependent, turn-on fluorescent protein sensor for chloride. ChemBioChem. 20, 1759–1765.

(24) Tutol, J. N., Ong, W. S. Y., Phelps, S. M., Peng, W., Goenawan, H., and Dodani, S. C. (2024) Engineering the ChlorON series: turn-on fluorescent protein sensors for imaging labile chloride in living cells. ACS Cent. Sci. 10, 77–86.

(25) Wang, Y., Xue, P., Cao, M., Yu, T., Lane, S. T., and Zhao, H. (2021) Directed evolution: methodologies and applications. Chem. Rev. 121, 12384–12444.

(26) Luo, W.-X., Cheng, T., Guan, B.-Q., Li, S.-W., Miao, J., Zhang, J., and Xia, N.-S. (2006) Variants of green fluorescent protein GFPxm. Mar. Biotechnol. 8, 560–566.

(27) Griesbeck, O., Baird, G. S., Campbell, R. E., Zacharias, D. A., and Tsien, R. Y. (2001) Reducing the environmental sensitivity of yellow fluorescent protein: mechanism and applications. J. Biol. Chem. 276, 29188–29194.

(28) Marcus, Y. (1994) A simple empirical model describing the thermodynamics of hydration of ions of widely varying charges, sizes, and shapes. Biophys. Chem. 51, 111–127.

(29) Ponten, J., and Saksela, E. (1967) Two established in vitro cell lines from human mesenchymal tumours. Int. J. Cancer. 2, 434–447.

(30) Weisbart, E., Kumar, A., Arevalo, J., Carpenter, A. E., Cimini, B. A., and Singh, S. (2024) Cell painting gallery: an open resource for image-based profiling. ArXiv. arXiv:2402.02203v1.

(31) Niforou, K. N., Anagnostopoulos, A. K., Vougas, K., Kittas, C., Gorgoulis, V. G., and Tsangaris, G. T. (2008) The proteome profile of the human osteosarcoma U2OS cell line. Cancer Genomics. 5, 63–78.

(32) Beck, M., Schmidt, A., Malmstroem, J., Claassen, M., Ori, A., Szymborska, A., Herzog, F., Rinner, O., Ellenberg, J., and Aebersold, R. (2011) The quantitative proteome of a human cell line. Mol. Syst. Biol. 7, 549.

(33) Rink, T. J., Tsien, R. Y., and Pozzan, T. (1982) Cytoplasmic pH and free Mg^2^+ in lymphocytes. J. Cell Biol. 95, 189–196.

(34) Sarkisyan, K. S., Goryashchenko, A. S., Lidsky, P. V., Gorbachev, D. A., Bozhanova, N. G., Gorokhovatsky, A. Y., Pereverzeva, A. R., Ryumina, A. P., Zherdeva, V. V., Savitsky, A. P., Solntsev, K. M., Bommarius, A. S., Sharonov, G. V., Lindquist, J. R., Drobizhev, M., Hughes, T. E., Rebane, A., Lukyanov, K. A., and Mishin, A. S. (2015) Green fluorescent protein with anionic tryptophan-based chromophore and long fluorescence lifetime. Biophys. J. 109, 380–389.

(35) Nasu, Y., Shen, Y., Kramer, L., and Campbell, R. E. (2021) Structure- and mechanism-guided design of single fluorescent protein-based biosensors. Nat. Chem. Biol. 17, 509–518.

(36) Wachter, R. M., Elsliger, M.-A., Kallio, K., Hanson, G. T., and Remington, S. J. (1998) Structural basis of spectral shifts in the yellow-emission variants of green fluorescent protein. Structure. 6, 1267–1277.

(37) Ong, W. S. Y., Ji, K., Pathiranage, V., Maydew, C., Baek, K., Villones, R. L. E., Meloni, G., Walker, A. R., and Dodani, S. C. (2023) Rational design of the β-bulge gate in a green fluorescent protein accelerates the kinetics of sulfate sensing. Angew. Chem. Int. Ed. Engl. 62, e202302304.

(38) Hendel, S. J., and Shoulders, M. D. (2021) Directed evolution in mammalian cells. Nat. Methods. 18, 346–357.

(39) Johnston, K. E., Fannjiang, C., Wittmann, B. J., Hie, B. L., Yang, K. K., and Wu, Z. (2023) Machine learning for protein engineering. ArXiv, arXiv:2305.16634v1.

