## Supporting Information for "Directed Evolution of a Genetically Encoded Indicator for Chloride"

### Methods

**General.** All chemicals, reagents, and supplies were purchased from Fisher Scientific, Sigma-Aldrich, or USA Scientific, except where noted.

**Multiple sequence alignment, percent identity matrix, and homology models.** Protein sequences were aligned using Clustal Omega.[S1] Homology models were generated using MODELLER V10.4.[S2] For GFPxm163 and OFPxm, the X-ray crystal structure of YFP-H148Q (PDB ID: 1F09) was used as the template.[S3] For mBeRFP, the X-ray crystal structure of mKate2 (PDB ID: 3SVN) was used as the template.[S4] PyMol 2.5.0 was used to visualize and render images of the crystal structures and homology models (Figure 1; Figure S1, S2, S11, S33–S37).

**Bacterial plasmid construction.** For expression in *Escherichia coli*, the gene encoding OFPxm (UniProt ID: Q8WTC5) was codon optimized and cloned into the pET-28a(+)-TEV vector between the NdeI and BamHI restriction sites to generate a N-terminal polyhistidine-tagged protein (GenScript, Figure S4). For the error-prone libraries, the gene encoding the mutated OFPxm was cloned between the same restriction sites (Figure S12).

**Protein expression, purification, and characterization.** The protein expression and purification methods were adapted from our previous report on GFPxm163 without any modification (Figure S5, S13).[S5] The protein characterization was carried out at room temperature (24–26 °C) following the methods for GFPxm163 with the following modifications.[S5] Depending on the protein and batch, the protein concentration used for characterization ranged from 2–6 μM and is specified in the figure captions. The chloride-bound p$K_a$ was measured in the presence of 100 mM $Cl^-$ instead of 512 mM $Cl^-$ [S5]. The pH titrations were carried out from pH 6.5–7.5 instead of pH 6–7.[S5] For the apparent dissociation constant ($K_d$) determination, the final anion concentrations used in the titrations were 0, 0.5, 1, 2, 4, 8, 16, 32, 64, and 128 mM. At each anion concentration, the fluorescence intensity at 524 nm (F) was first normalized ($F_n$). The minimal fluorescence intensity in the presence of anions ($F_{min}$) was extrapolated to 0 for all anions tested at all pH conditions, and the maximum fluorescence intensity ($F_{max}$) is F with 0 mM anion. The $F_n$ was fitted into a single binding site model to determine the $K_d$ using the equation below (KaleidaGraph v. 4.5.3, Synergy). All data is shown in Figure 2 in the Main Text and Figure S6, S7, S14–S25, and Table S4 in the Supporting Information.

$$F = (F_{max} - F_{min}) \times \left(1 - \frac{[anion]}{K_d + [anion]}\right) + F_{min} = F_{max} \times (1 - \frac{[anion]}{K_d + [anion]})$$

$$F_n = \frac{F}{F_{max}} = \frac{K_d}{K_d + [anion]}$$

**Cloning.** The commercial pET-28a(+)-TEV-OFPxm plasmid was used as the starting template for the construction of the first error-prone library. The error-prone polymerase chain reaction (EP-PCR) and high-fidelity PCR were carried out in parallel with two sets of complementary primers to generate the OFPxm insert with mutations and unmutated vector backbone, respectively (Table S1). Taq polymerase (New England Biolabs) was used with an addition of $MnCl_2$ to amplify the OFPxm insert with mutations. Mutagenesis was carried out with 50, 100, 200, 300 µM $MnCl_2$ to determine that 200 µM was optimal. The mutation rate was confirmed to be 3 base pairs per 1,000 base pairs by sequencing 3 random variants from the same library (Eurofins). Phusion Hot Start Flex (New England Biolabs) was used to amplify the pET-28a(+)-TEV backbone. The detailed reaction components and conditions for the OFPxm insert and vector backbone are shown in Table S2 and S3, respectively.

The PCR product was incubated with 1 µL of Dpn1 at 37 °C for 2 h to remove any template DNA. The DNA fragments were separated with agarose gel (1% agarose in 1X Tris-Acetate-EDTA buffer) electrophoresis and extracted with a Zymoclean Gel DNA Recovery Kit (Zymo Research). The concentrations of the purified DNA fragments were determined using a NanoDrop Lite Spectrophotometer (Fisher Scientific). For plasmid assembly, a total of 100 ng of the DNA (3:1 ratio of the insert to the backbone) was diluted with autoclaved water to a volume of 10 µL and incubated with 10 µL of 2X HiFi Assembly Mater Mix (New England Biolabs) at 50 °C for 1 h. The reaction product was purified with a DNA Clean & Concentrator Kit (Zymo Research) and stored at -20 °C. All steps above were repeated for each round with a new parent plasmid from the previous round. The library expression and screening methods are described below.

**Error-prone library expression and screening.** For each round, the parent and error-prone library prepared above in the *Cloning section* was used to transform *E. cloni* EXPRESS BL21 (DE3) competent cells (Lucigen) by electroporation (MicroPulser Electroporator, Bio-Rad Laboratories). The cells were recovered at 37 °C for 30 min with shaking at 250 rpm in super optimal broth with catabolite repression or SOC media, plated onto LB (Luria Broth, Research Products International) agar plates with 50 µg/mL kanamycin, and incubated at 37 °C for 18 h. Prior to the first round of library expression and screening, four library plates, each using a different concentration of $MnCl_2$, were expressed and screened to optimize the $MnCl_2$ concentration for a moderate mutation rate. For each round of directed evolution, ~1,800 variants were expressed and screened as follows. For each library plate, 8 single colonies of the parent and 86 colonies from the mutated library were picked into 500 µL of LB with 50 µg/mL kanamycin in a 96-deep-well plate format. Two wells were left empty to serve as a negative control. All plates were sealed with an EasyApp microporous film (USA Scientific) and incubated at 37 °C for 16 h with shaking at 250 rpm. The next day, 50 µL of the overnight culture was transferred into another 96-deep-well plate filled with 950 µL of LB with 50 µg/mL kanamycin by a liquid handler (Biomek $NX^P$, Beckman Coulter) and further incubated at 37 °C for 2 h with shaking at 250 rpm. Following this, 50 µL of LB with 50 µg/mL kanamycin and 21 mM isopropyl beta-D-thiogalactopyranoside (IPTG, Gold Biotechnology) was added to each well to induce protein expression. After incubation at 37 °C for 18 h with shaking at 250 rpm, the cells were harvested by centrifugation at 3,000*g* (5810 R, Eppendorf) for 15 min at 4 °C. The resulting cell pellets were stored at -20 °C.

For plate screening, the frozen cell pellets were thawed for 20 min at room temperature and then resuspended in 50 µL of B-PER (Bacterial Protein Extraction Reagent) by briefly agitating on a vortexer (Vortex 2, IKA) with a microtiter plate attachment. The lysis proceeded for 15 min at room temperature. Following this, 500 µL of 50 mM sodium phosphate buffer at pH 7 was added to each well and mixed with agitation on a vortexer. After centrifugation at 3,000*g* for 15 min at 20 °C, 175 µL of the supernatant from each well was transferred into a 96-well microtiter plate using the liquid handler for analysis using a microplate reader (Spark, Tecan) at room temperature (24–26 °C). Prior to scanning on the microplate reader, the plate was subjected to double orbital shaking for 10 s with an amplitude of 2.5. For each well, excitation was provided at 480 nm (5 nm bandwidth), and the emission intensity was collected from 500–540 nm (10 nm step size, 5 nm bandwidth, 30 flashes). Then, 25 µL of 50 mM sodium phosphate buffer at pH 7 with 800 mM NaBr was added into each well with a multichannel pipet to a final concentration of 100 mM NaBr for the first round. For the second and third rounds, 400 mM NaBr or NaCl stock solutions were used (50 mM final concentration), respectively. Prior to re-scanning, the plate was subjected to shaking again with the same settings. The wells with a low fluorescence signal-to-noise ratio (relative fluorescence units < 500) were removed from the analysis. For each well, the fold-change was calculated by dividing the emission intensity at 520 nm in the presence of the anion by the emission intensity at 520 nm in the absence of the anion. The resulting fold-changes were sorted and plotted corresponding to the well identity to generate the ROF curve. For each plate, the fold-change of all parent wells in the plate were averaged to calculate the mean and the standard deviation. A threshold of three standard deviations above and below the mean for the parent fold change was used to select improved variants for further validation. The fold-changes of all the parent wells across the plate were averaged to calculate the mean and standard deviation across the entire library for each round (Figure S8, S9).

The improved variants from each round were re-streaked from the overnight source plate onto LB agar plates with 50 µg/mL kanamycin and incubated at 37 °C for 16 h. Three single colonies of each variant and retransformed parent were picked and inoculated into 1 mL of LB with 50 µg/mL kanamycin in 5 mL culture tubes. After incubation at 37 °C for 2 h with shaking at 250 rpm, 1 µL of 1 M IPTG was added to induce protein expression at 37 °C for 18 h with shaking at 250 rpm. The following day, each expression culture was transferred into a 1.5 mL microcentrifuge tube and collected by centrifugation at 18,000*g* (5430 R, Eppendorf) for 10 min at 4 °C. The resulting cell pellet was resuspended in 50 µL of B-PER by agitation on a vortexer and maintained at room temperature for lysis. After 15 min, 1 mL of 50 mM sodium phosphate buffer at pH 7 was added. The cell suspension was clarified by centrifugation at 18,000*g* for 10 min at 20 °C. Next, 175 µL of the lysate was combined with 25 µL of 50 mM sodium phosphate buffer at pH 7 with 0, 200, 400, and 800 mM NaBr in the microtiter plate to a final concentration of 0, 25, 50, and 100 mM NaBr, respectively. For each well, the absorbance was collected from 350–600 nm (5 nm step size), excitation was provided at 480 nm (5 nm bandwidth), and the emission intensity was collected from 500–600 nm (5 nm step size, 5 nm bandwidth, 30 flashes) (Figure S10). The fold-change was calculated by dividing the emission intensity at 520 nm in the presence of the anion by the emission intensity at 520 nm in the absence of the anion. The data from all three replicates was averaged to calculate the mean and standard deviation. The plasmid DNA from each variant with a statistically higher degree of quenching than the parent ($p < 0.001$) was extracted using the QIAprep Spin Miniprep Kit. Following this, the mutations were identified by Sanger Sequencing (Eurofins) (Figure 1; Figure S11).

**Mammalian plasmid construction and preparation.** For expression in mammalian cells, the gene encoding ChlorOFF was codon optimized and cloned into the pcDNA3.1(+) vector between the BamHI and EcoRI restriction sites with a C-terminal stop codon (GenScript, Figure S26). The pcDNA3.1(+)-ChlorOFF plasmid was used to transform One Shot TOP10 Chemically Competent *E. coli* by heat shock according to the manufacturer's instructions and recovered in SOC at 37 °C for 1 h with shaking at 250 RPM. Fifty microliters of the recovered cell culture were plated on a LB agar plate with 100 µg/mL ampicillin and incubated at 37 °C for 18 h (Innova42, New Brunswick). A single colony was inoculated in 100 mL of LB with 100 µg/mL ampicillin and incubated at 37 °C for 16 h with shaking at 250 rpm. The plasmid was extracted using the QIAprep Spin Maxiprep Kit (Qiagen) and stored at –20 °C.

**U-2 OS cell culture and ChlorOFF transfection, selection, sorting, and enrichment.** The U-2 OS (HTB-96) cells were purchased from the American Type Culture Collection.[S6] The cells were cultured in flasks using McCoy's 5A media supplemented with 10% fetal bovine serum (FBS) and 1% Penicillin-Streptomycin (Pen-Strep, 100 µg/mL) at 37 °C, 5% $CO_2$. Once flasks reached 90–100% confluency, cells were washed with Phosphate Buffered Saline (PBS), treated with 0.05% trypsin-EDTA, and incubated for ~5–7 min at 37 °C, 5% $CO_2$. The trypsin reaction was quenched with McCoy's 5A media supplemented with 10% FBS and 1% Pen-Strep, and the cells were harvested via centrifugation at 200*g* for 5 min (5702, Eppendorf) and resuspended in 3–5 mL media. A hemocytometer was used to determine the cell count, and ~$6 \times 10^5$ cells in 2 mL complete media were seeded per well into 5 wells of a 6-well plate for transfection the next day. The remaining 6th well was seeded with ~$3 \times 10^5$ cells to serve as a negative control during the fluorescence sorting. The plate was incubated overnight at 37 °C, 5% $CO_2$, until the wells were at least 70% confluent. The following day, a solution of 1.25 mL OptiMEM Reduced Serum, 11.25 µL of 1 µg/mL pcDNA3.1(+)-ChlorOFF, 22.5 µL P3000, and 18.75 µL Lipofectamine 3000 was prepared in a 1.5 mL tube, lightly agitated by a vortexer, and incubated at room temperature for at least 30 min. Then, 250 µL of the lipid-pcDNA mixture was added in a dropwise fashion to each well, and the plate was incubated for 3 days at 37 °C, 5% $CO_2$.

On the day of sorting, each well was washed with 2 mL PBS, treated with 1 mL 0.05% trypsin-EDTA, incubated for ~5–7 min at 37 °C, 5% $CO_2$, and the trypsinization was quenched with 2 mL McCoy's 5A media supplemented with 10% FBS. The transfected wells were combined in a 14 mL conical centrifuge tube and harvested via centrifugation at 300*g* for 5 min. The cell pellet was then washed with 137 mM sodium gluconate (Gluc) buffer containing 8.1 mM $Na_2HPO_4$, 1.5 mM $KH_2PO_4$, 2.7 mM KGluc, 0.7 mM $CaGluc_2$, 1.1 mM $MgGluc_2$, and 2% FBS and harvested again at 200*g* for 5 min. The supernatant was aspirated, and the cell pellet was resuspended in 700 µL 137 mM NaGluc buffer supplemented with 2% FBS, incubated at 37 °C for 30 min, and transferred to a 5 mL flow cytometry tube with a strainer cap to reduce cell clumping. Using the FACSAria Fusion Flow Cytometer (BD Biosciences) in the UT Dallas Flow Cytometry Core, the GFP filter was used to sort for cells expressing ChlorOFF. Non-transfected cells were used to draw gates and exclude the cell population. All ChlorOFF-positive cells were collected in a 14 mL conical centrifuge tube containing McCoy's 5A media supplemented with 10% FBS, harvested via centrifugation at 300*g* for 5 min, and resuspended in McCoy's 5A media supplemented with 10% FBS and 500 µg/mL G148. The cells were then transferred to a T25 flask and incubated at 37 °C, 5% $CO_2$. Once the flask was 100% confluent, the sorting process was repeated to generate an enriched U-2 OS cell population expressing ChlorOFF.

**Anion exchange microscopy assays with ChlorOFF.** The U-2 OS ChlorOFF cells were cultured and passaged in McCoy's 5A media supplemented with 10% FBS and 500 µg/mL G148 as described above. For fluorescence microscopy assays, ~7–8 x $10^5$ cells expressing ChlorOFF were plated onto 35 mm dishes with a 14 mm micro-well and #1.5 glass-like polymer coverslip (Cellvis) in 2 mL media and incubated overnight at 37 °C, 5% $CO_2$. On the day of imaging, each dish was washed twice with 2 mL of a modified PBS buffer at pH 7.4 containing 137 mM NaCl, 8.1 mM $Na_2HPO_4$, 1.5 mM $KH_2PO_4$, 2.7 mM KCl, 0.7 mM $CaCl_2$, 1.1 mM $MgCl_2$, and 10 mM glucose or 137 mM NaGluc, 8.1 mM $Na_2HPO_4$, 1.5 mM $KH_2PO_4$, 2.7 mM KGluc, 0.7 mM $CaGluc_2$, 1.1 mM $MgGluc_2$, and 10 mM glucose. The osmolarity of all buffers were measured at ~280 mOsm (VAPRO, ELITechGroup). Each dish was incubated on stage or in an incubator for 30 min at 37 °C in the modified PBS buffer. All differential interference contrast (DIC) and fluorescence images were acquired on an inverted fluorescence microscope (IX83, Olympus) equipped with a light engine (Spectra X, Lumencor) at 37 °C. A 20X air objective with a numerical aperture of 0.7 and a working distance of 1.6 mm and an EGFP filter set (Chroma) were used. Excitation was provided at 488 nm with 25% LED power. The exposure time ranged from 175–275 ms. The camera resolution was set to 512 x 512 pixels with 2 x 2 binning. The coordinates for three different fields were recorded with Z-drift compensation (ZDC) for each dish.

The following methods were adapted from our previous study.[S7] Images were acquired every 30 s for 5 min before the imaging solution was exchanged with 8 mL of a modified PBS buffer at pH 7.4 containing 137 mM NaCl (or 68.5 mM NaI/68.5 mM NaCl), 8.1 mM $Na_2HPO_4$, 1.5 mM $KH_2PO_4$, 2.7 mM KCl, 0.7 mM $CaCl_2$, 1.1 mM $MgCl_2$, and 10 mM glucose or 137 mM NaGluc, 8.1 mM $Na_2HPO_4$, 1.5 mM $KH_2PO_4$, 2.7 mM KGluc, 0.7 mM $CaGluc_2$, 1.1 mM $MgGluc_2$, and 10 mM glucose at a rate of 4 mL/min for 2 min using an automated perfusion system (Tokai Hit). Images were then acquired every 30 s for 25 min during perfusion.

**U-2 OS staining and anion exchange microscopy assays using a pH dye.** The following methods were adapted from our previous study.[S7] The pH indicator dye 2',7'-bis-(2-carboxyethyl)-5-(and-6)-carboxyfluorescein, acetoxymethyl ester (BCECF-AM, Invitrogen) was dissolved in anhydrous DMSO and stored as 10 mM aliquots at -20 °C. On the day of imaging, BCECF-AM was diluted to 5 mM with anhydrous DMSO. U-2 OS cells were plated onto 35 mm dishes as described above at a seeding density of ~6–7 × $10^5$ cells and incubated at 37 °C, 5% $CO_2$. The next day, the cells were washed with 2 mL DMEM containing 1% Pen-Strep and stained with 5 μM BCECF-AM in DMEM containing 1% Pen-Strep and incubated for 1 h at 37 °C, 5% $CO_2$. The anion exchange assay was carried out at 37 °C as described above with the following modifications. Images were acquired every 1 min, and a custom filter set for BCECF emission at 540 nm (Chroma) was used for the excitation at 436 nm (cyan) and 495 nm (red) with the exposure time set to 80 ms. For each dish, the coordinates were recorded for two different fields with ZDC. At the end of the experiment, the imaging solution was manually exchanged with a pH clamping solution containing 120 mM KGluc, 20 mM NaGluc, 8.1 mM $Na_2HPO_4$, 1.5 mM $KH_2PO_4$, 0.7 mM $CaGluc_2$, 1.1 mM $MgGluc_2$, 5 μM valinomycin, and 5 μM nigericin at pH 8. The dish was incubated at 37 °C for an additional 15 min before images were reacquired.

**Image process and analysis.** The Fiji is Just ImageJ software (Fiji v2.0) was used as described in our previous study with the following modifications.[S5,S8] All fluorescence channels were split and aligned using the StackReg Translation function. From these, the background was subtracted (size 75). A mask was generated using the maximum intensity Z-projection for ChlorOFF and the cyan excitation channel for BCECF by adjusting the minimum default threshold to 100 for ChlorOFF and 750 for BCECF (Figure S27, S29). Watershed separation was then used to further delineate the mask between cells or cell clusters, and ROIs were selected using the Analyze Particles function (Figure S27, S29). The ROIs were transferred to each background-subtracted fluorescence stack. From these, the median intensity was measured for each ROI. Outliers in the ChlorOFF florescence responses (F) were determined using Microsoft Excel to calculate the Interquartile Range (IQR) which is the difference between Quartile 3 (Q3) and Quartile 1 (Q1) for each data set. The upper and lower ranges were calculated using the following equations, and data that fell outside of these ranges were considered outliers and excluded from our analyses:

$$\text{Lower range: } Q1 - (IQR*1.5)$$
$$\text{Upper range: } Q3 + (IQR*1.5)$$

For each experiment, the average median intensity for three biological replicates with standard deviation is reported (Figure 3; Figure S28, S30–S32, Movie S1–S6).

### *Figures and Tables*

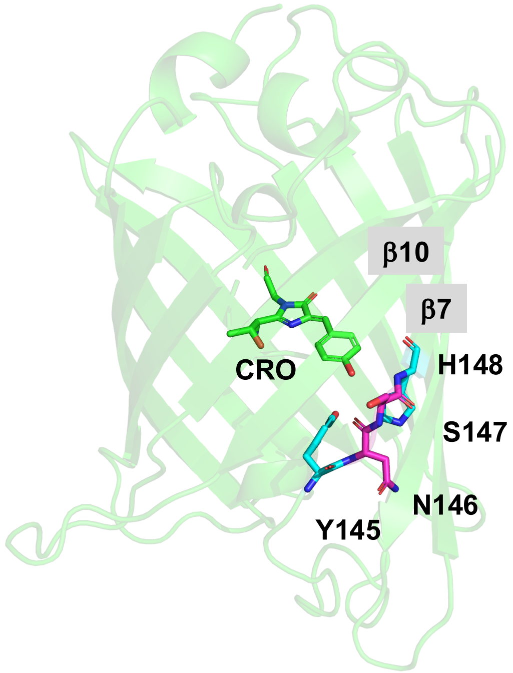


**Figure S1.** X-ray crystal structure of the green fluorescent protein from the jellyfish *Aequorea victoria* (avGFP, PDB ID: 1EMA) with the chromophore (green sticks), gate post (cyan sticks), and b-bulge (magenta sticks). The β7 and β10 strands are labeled for reference. Abbreviation: CRO, chromophore.

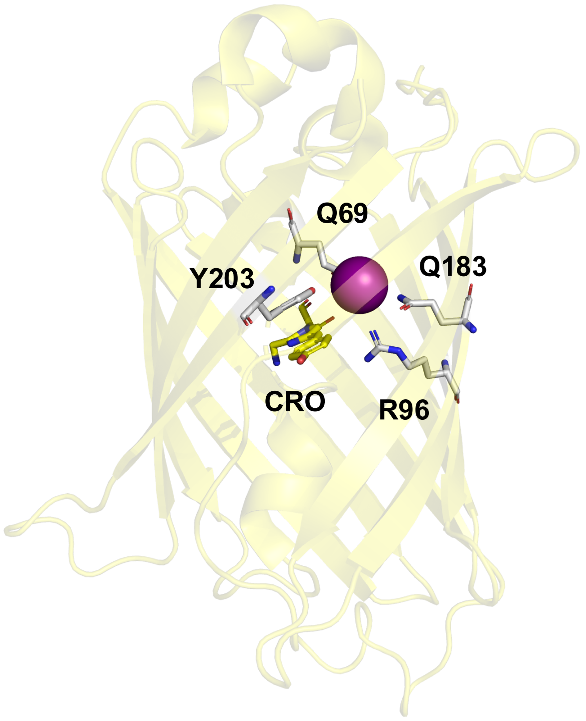


**Figure S2.** X-ray crystal structure of the anion-sensitive avGFP variant avYFP-H148Q (PDB ID: 1F09) with the chromophore (yellow sticks), binding pocket residues (gray sticks), and iodide (purple sphere). Abbreviation: CRO, chromophore.

**A**

```
CLUSTAL O(1.2.4) multiple sequence alignment

YFP-H148Q       MSKGEELFTGVVPILVELDGDVNGHKFSVSGEGEGDATYGKLTLKFICTTGKLPVPWPTL 60
GFPxm163        MSKGEELFTGIVPVLIELDGDVHGHKFSVRGEGEGDADYGKLEIKFICTTGKLPVPWPTL 60
OFPxm           MSKGEELFTGVVPILVELDGDVHGHKFSVRGEGEGDADYGKLEIKFICTTGKLPVPWPTL 60
                **********:**:*:******:****** ******* **** :****************

YFP-H148Q       VTTFGYGLQCFARYPDHMKRHDFFKSAMPEGYVQERTIFFKDDGNYKTRAEVKFEGDTLV 120
GFPxm163        VTTLGYGIQCFARYPEHMKMNDFFKSAMPEGYIQERTIFFQDDGKYKTRGEVKFEGDTLV 120
OFPxm           VTTLGYGILCFARYPEHMKMNDFFKSAMPEGYIQERTIFFQDDGKYKTRGEVKFEGDTLV 120
                ***:***: ******:*** :***********:*******:***:****.**********

YFP-H148Q       NRIELKGIDFKEDGNILGHKLEYNYNSQNVYIMADKQKNGIKVNFKIRHNIEDGSVQLAD 180
GFPxm163        NRIELKGMDFKEDGNILGHKLEYNFNSHNVYIMPDKANNGLKVNFKIRHNIEGGGVQLAD 180
OFPxm           NRIELKGMDFKEDGNILGHKLEYNFNSHNVYIMPDKANNGLKVNFKIRHNIEGGGVQLAD 180
                *******:****************:**:***** ** :**:***********.*.*****

YFP-H148Q       HYQQNTPIGDGPVLLPDNHYLSYQSALSKDPNEKRDHMVLLEFVTAAGITHGMDELYK 238
GFPxm163        HYQTNVPLGDGPVLIPINHYLSYQTAISKDRNETRDHMVFLEFFSACGHTHGMDELYK 238
OFPxm           HYQTNVPLGDGPVLIPINHYLSYQTAISKDRNETRDHMVFLEFFSACGHTHGMDELYK 238
                *** *.*:******:* *******:*:*** **.*****:***.:*.* *********
```

**B**

```
Percent Identity  Matrix - created by Clustal2.1

     1: YFP-H148Q    100.00   83.19   84.03
     2: GFPxm163      83.19  100.00   98.32
     3: OFPxm         84.03   98.32  100.00
```

**Figure S3.** (A) Multiple sequence alignment of avYFP-H148Q, GFPxm163, and OFPxm. The chromophore is highlighted in green, and the anion binding pocket residues are highlighted in cyan. (B) Percent identity matrix of avYFP-H148, GFPxm163, and OFPxm.

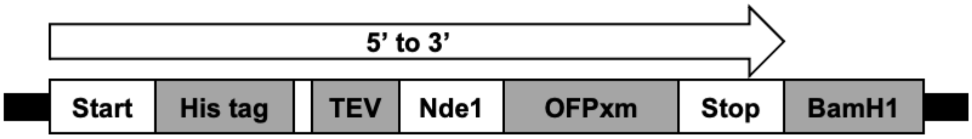


ATGGGCAGCAGCCATCATCATCATCATCACAGCAGCGGCGAGAATCTTTATTTTCAGGGCC
ATATGAGCAAGGGCGAGGAACTGTTCACCGGCGTGGTTCCGATCCTGGTGGAGCTGGAC
GGTGATGTTCACGGCCACAAATTTAGCGTTCGTGGCGAGGGTGAAGGTGATGCGGATTAC
GGCAAGCTGGAAATCAAATTCATTTGCACCACCGGTAAACTGCCGGTTCCGTGGCCGACC
CTGGTTACCACCCTGGGTTACGGCATTCTGTGCTTTGCGCGTTATCCGGAGCACATGAAGA
TGAACGACTTCTTTAAAAGCGCGATGCCGGAGGGTTACATCCAGGAACGTACCATTTTCTT
TCAAGACGATGGCAAGTACAAGACCCGTGGCGAGGTGAAATTCGAAGGTGATACCCTGGT
TAACCGTATCGAGCTGAAGGGCATGGACTTCAAAGAAGATGGTAACATTCTGGGCCACAA
GCTGGAGTACAACTTTAACAGCCACAACGTGTATATCATGCCGGACAAAGCGAACAACGGT
CTGAAGGTTAACTTTAAAATCCGTCACAACATTGAAGGTGGCGGTGTGCAGCTGGCGGAC
CACTACCAAACCAACGTGCCGCTGGGTGATGGCCCGGTTCTGATCCCGATTAACCACTAC
CTGAGCTATCAGACCGCGATTAGCAAGGACCGTAACGAGACCCGTGATCACATGGTGTTC
CTGGAATTCTTTAGCGCGTGCGGTCACACCCACGGCATGGATGAACTGTATAAATAAGGAT
CC

MGSSHHHHHHSSGENLYFQGHMSKGEELFTGVVPILVELDGDVHGHKFSVRGEGEGDADYG
KLEIKFICTTGKLPVPWPTLVTTLGYGILCFARYPEHMKMNDFFKSAMPEGYIQERTIFFQDDGK
YKTRGEVKFEGDTLVNRIELKGMDFKEDGNILGHKLEYNFNSHNVYIMPDKANNGLKVNFKIRH
NIEGGGVQLADHYQTNVPLGDGPVLIPINHYLSYQTAISKDRNETRDHMVFLEFFSACGHTHGM
DELYK

**Figure S4.** The pET-28a(+)-TEV-OFPxm plasmid design expression in *Escherichia coli* (top panel), nucleotide sequence (middle panel), and amino acid sequence (bottom panel).

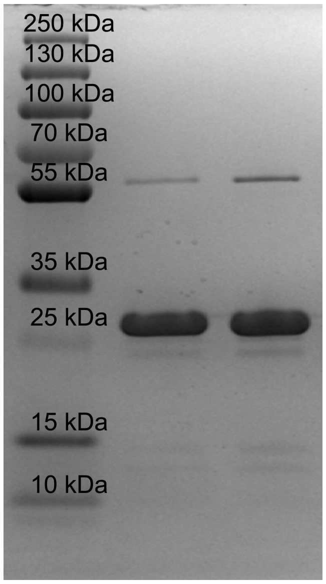


**Figure S5.** A representative Coomassie-stained SDS-PAGE gel of all OFPxm protein batches used for characterization in this study. The calculated molecular weight for OFPxm is ~29 kDa.

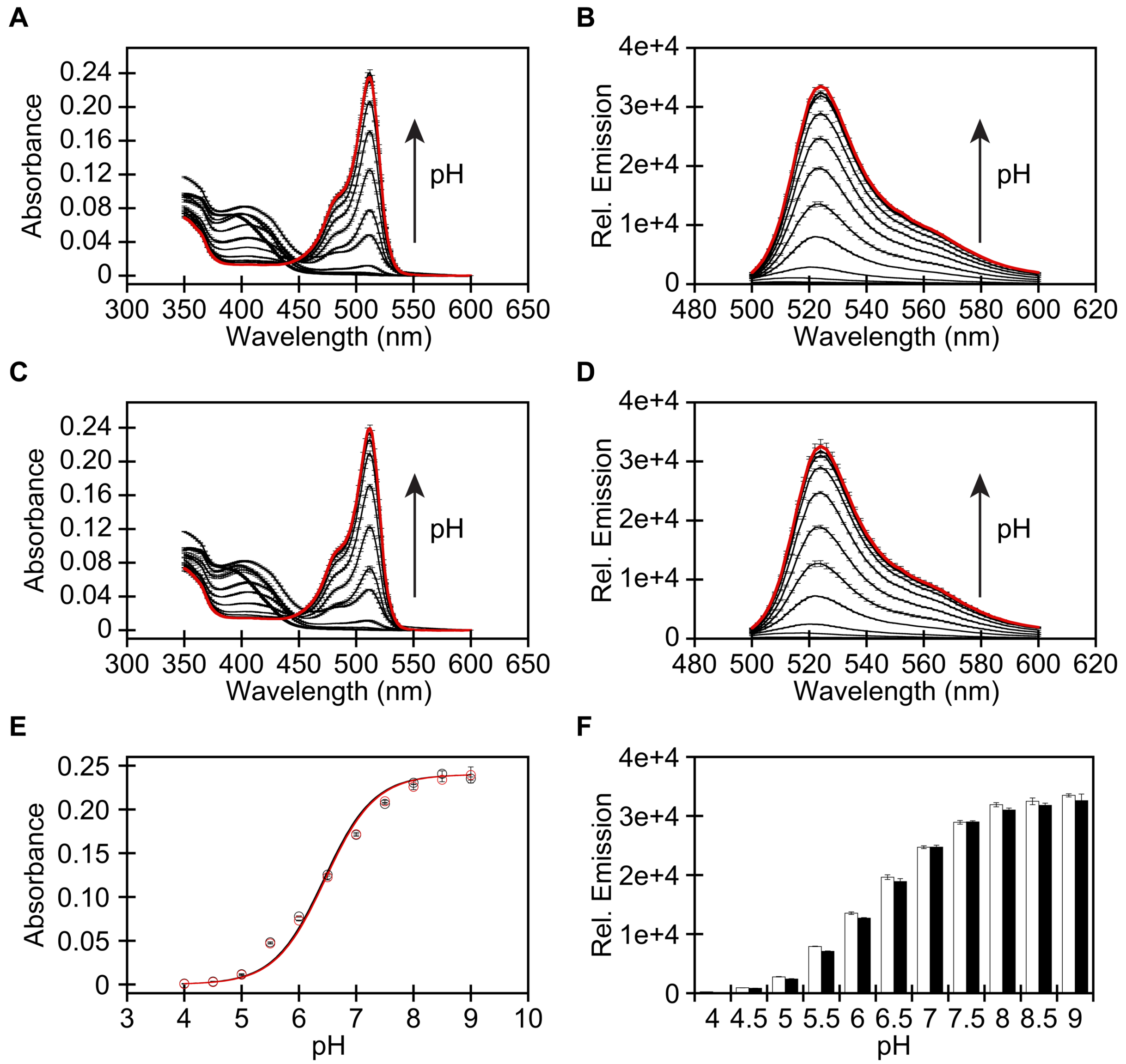


**Figure S6.** (A) Absorbance ($\lambda_{abs}$ = 350–600 nm) and (B) emission ($\lambda_{ex}$ = 480 nm, $\lambda_{em}$ = 500–600 nm) spectra of 6 μM OFPxm from pH 4 (bold black) to pH 9 (red). (C) Absorbance and (D) emission spectra of 6 μM OFPxm from pH 4 (bold black) to pH 9 (red) in the presence of 100 mM $Cl^-$. (E) Absorbance-dependent p$K_a$ curve of OFPxm with 0 mM (black) and 100 mM (red) $Cl^-$ at $\lambda_{abs}$ = 512 nm. The p$K_a$ values in the absence and presence of chloride are 6.44 ± 0.05 and 6.46 ± 0.05, respectively. (F) Emission response of OFPxm with 0 mM (white) and 100 mM (black) $Cl^-$ at $\lambda_{em}$ = 524 nm. Data is shown for one of two protein preparations (average with S.E.M.). Data for the second protein preparation is shown in Figure S7. The following buffers were used: 50 mM sodium acetate buffer for pH 4–5.5 or 50 mM sodium phosphate buffer for pH 6–9. Abbreviation: standard error of the mean, S.E.M.

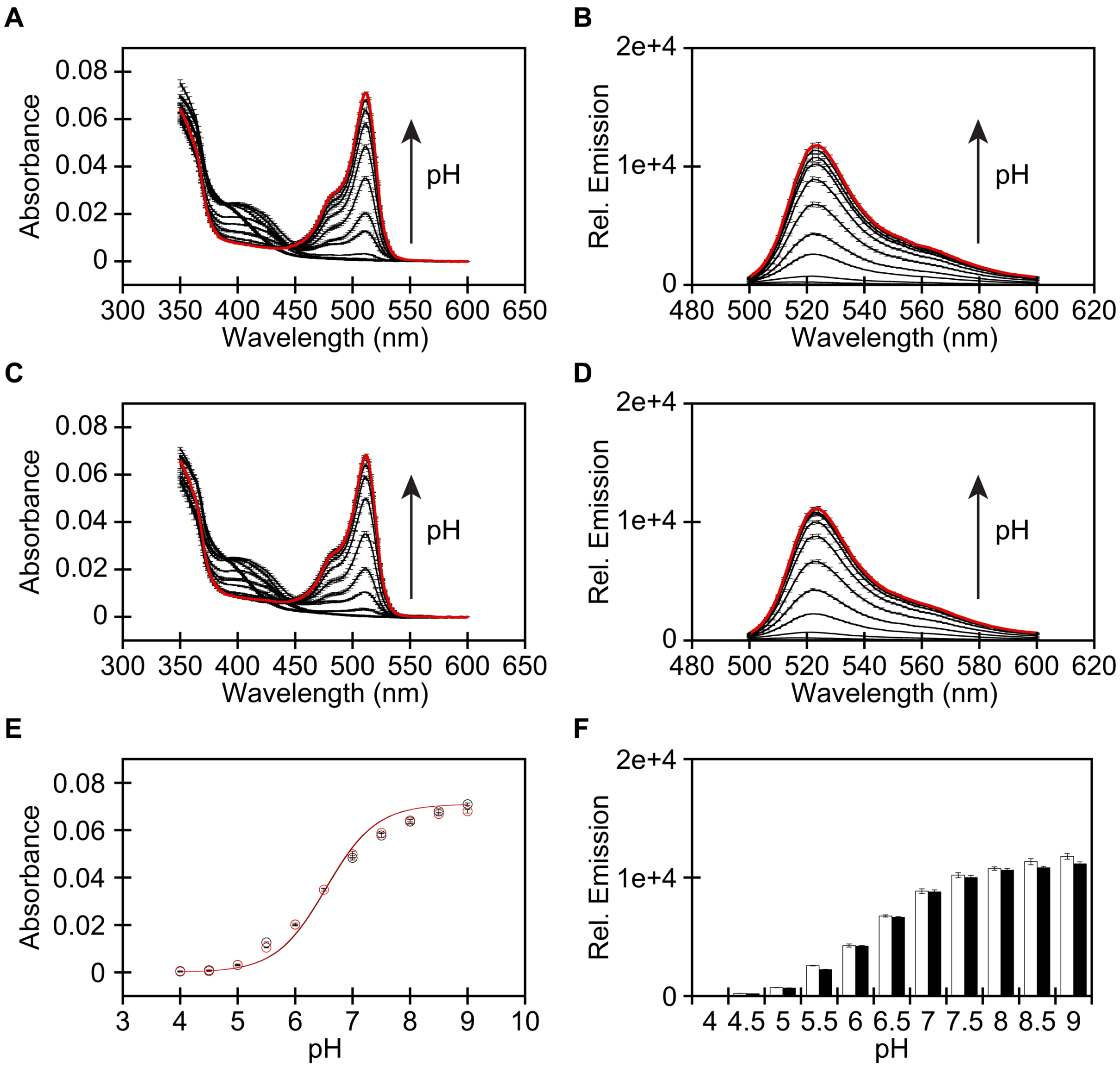


**Figure S7.** (A) Absorbance ($\lambda_{abs}$ = 350–600 nm) and (B) emission ($\lambda_{ex}$ = 480 nm, $\lambda_{em}$ = 500–600 nm) spectra of 2 μM OFPxm from pH 4 (bold black) to pH 9 (red). (C) Absorbance and (D) emission spectra of 2 μM OFPxm from pH 4 (bold black) to pH 9 (red) in the presence of 100 mM $Cl^-$. (E) Absorbance-dependent p$K_a$ curve of OFPxm with 0 mM (black) and 100 mM (red) $Cl^-$ at $\lambda_{abs}$ = 512 nm. The p$K_a$ values in the absence and presence of chloride are 6.54 ± 0.06 and 6.53 ± 0.05, respectively. (F) Emission response of OFPxm with 0 mM (white) and 100 mM (black) $Cl^-$ at $\lambda_{em}$ = 524 nm. Data is shown for one of two protein preparations (average with S.E.M.). Data for the second protein preparation is shown in Figure S6. The following buffers were used: 50 mM sodium acetate buffer for pH 4–5.5 or 50 mM sodium phosphate buffer for pH 6–9.

**Table S1.** Polymerase chain reaction (PCR) primers for the error-prone libraries.

| Primers | | Sequence (5' → 3') |
|---|---|---|
| OFPxm insert | Forward | GCGAGAATCTTTATTTTCAGGGCCATATG |
| | Reverse | CGGAGCTCGAATTCGGATCCTTA |
| pET-28a(+)-TEV backbone | Forward | TAAGGATCCGAATTCGAGCTCCG |
| | Reverse | CATATGGCCCTGAAAATAAAGATTCTCGC |

**Table S2.** PCR components (top) and thermocycler settings (bottom) for the mutated OFPxm insert with Taq polymerase.

| Components | Volume (μL) | | | |
|---|---|---|---|---|
| **10X Taq buffer** | 5 | 5 | 5 | 5 |
| **10 μM dNTPs** | 1 | 1 | 1 | 1 |
| **10 mM forward primer** | 1 | 1 | 1 | 1 |
| **10 mM reverse primer** | 1 | 1 | 1 | 1 |
| **10 ng/μL template DNA** | 2 | 2 | 2 | 2 |
| **Taq DNA polymerase** | 0.25 | 0.25 | 0.25 | 0.25 |
| **4 mM $MnCl_2$** | 0.63 | 1.25 | 2.5 | 3.75 |
| **Autoclaved water** | 39.12 | 38.5 | 37.25 | 36 |
| **Final [$MnCl_2$] (μM)** | 50 | 100 | 200 | 300 |

| Step | Temperature (°C) | Time (s) | cycle |
|---|---|---|---|
| **Initial denaturation** | 95 | 30 | 1 |
| **Denaturation Annealing Elongation** | 95 | 30 | 30 |
| | 55 | 30 | |
| | 68 | 60 | |
| **Final elongation** | 68 | 600 | 1 |
| **Hold** | 10 | ∞ | 1 |

**Table S3.** PCR components (top) and thermocycler settings (bottom) for the pET-28a(+)-TEV backbone with Phusion.

| Components | Volume (μL) |
|---|---|
| **Phusion Hot Start Flex 2X master mix** | 50 |
| **10 mM forward primer** | 2 |
| **10 mM reverse primer** | 2 |
| **10 ng/μL template DNA** | 2 |
| **Dimethyl sulfoxide** | 3 |
| **Autoclaved water** | 41 |

<table>
<tr><th>Step</th><th>Temperature (°C)</th><th>Time (s)</th><th>cycle</th></tr>
<tr><td>Initial denaturation</td><td>98</td><td>30</td><td>1</td></tr>
<tr><td rowspan="3">Denaturation<br>Annealing<br>Elongation</td><td>98</td><td>10</td><td rowspan="3">30</td></tr>
<tr><td>70</td><td>30</td></tr>
<tr><td>72</td><td>150</td></tr>
<tr><td>Final elongation</td><td>72</td><td>600</td><td>1</td></tr>
<tr><td>Hold</td><td>10</td><td>∞</td><td>1</td></tr>
</table>

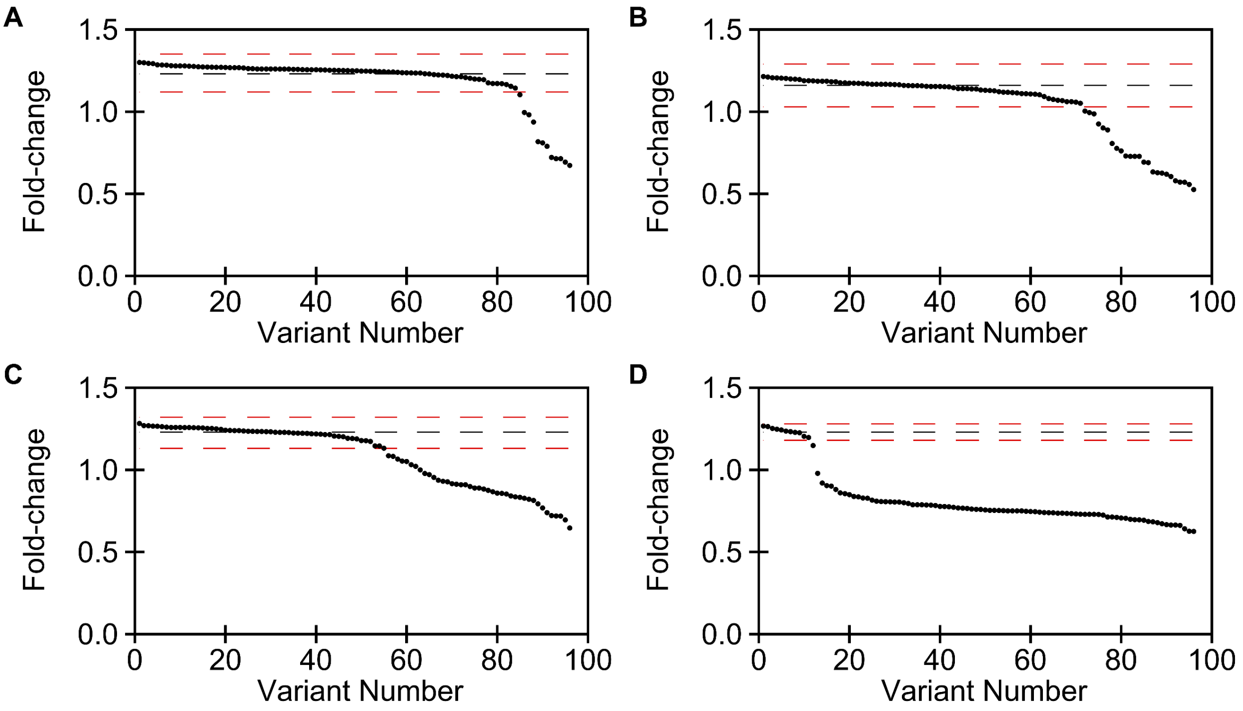


**Figure S8.** The retention of function curves for pilot libraries generated with (A) 50 μM, (B) 100 μM, (C) 200 μM, and (D) 300 μM $MnCl_2$ in the EP-PCR reaction. The averaged fold-change of the parent is marked as black dashed line, and the selection threshold is marked as red dashed line. Each dot represents one library variant.

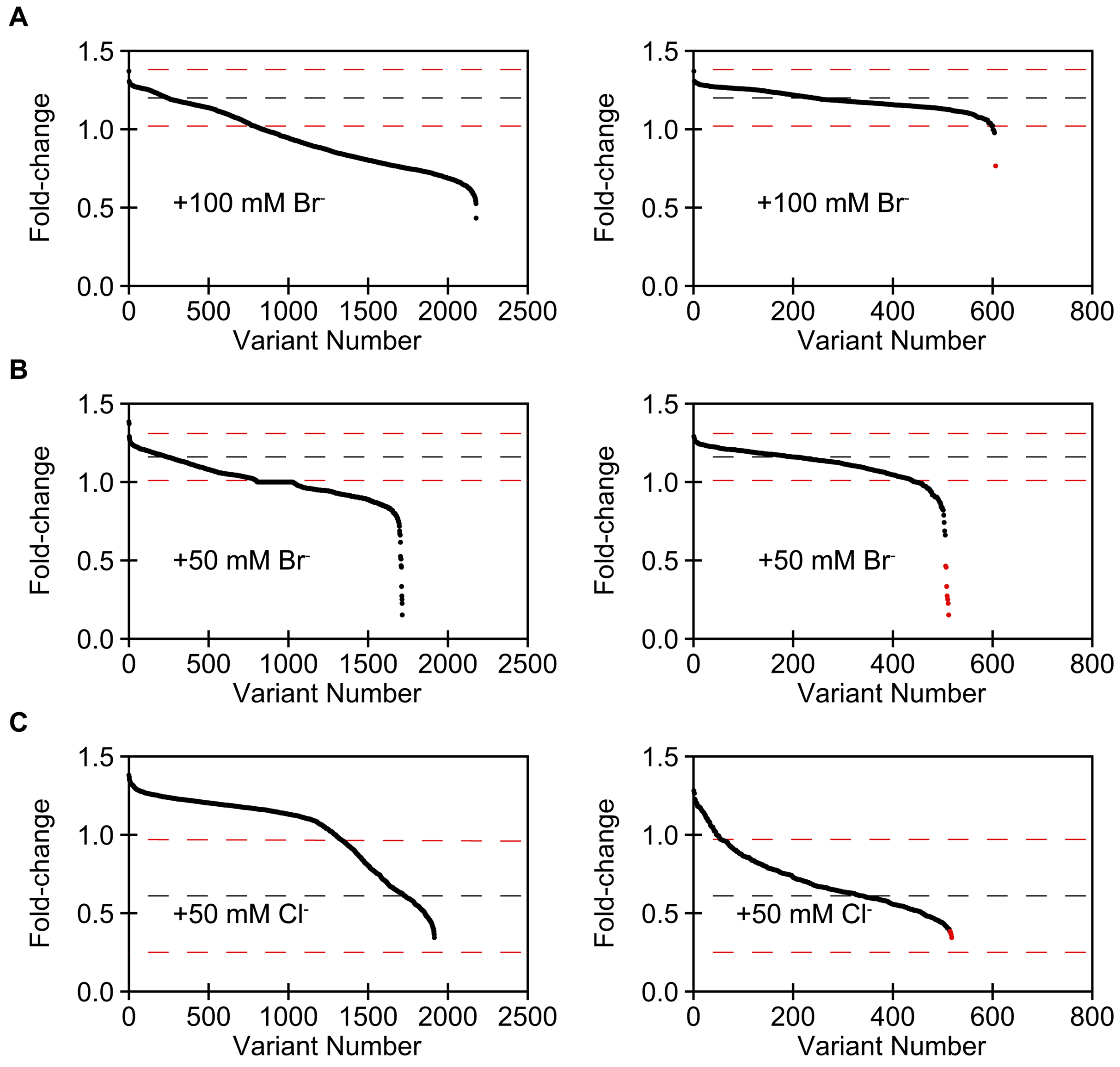


**Figure S9.** The calculated fold-change for all library variants before (left) and after (right) filtering the ones with fluorescent intensity < 500 for round (A) one, (B) two, and (C) three of the directed evolution. The averaged fold-change of the parent is marked as black dashed line, and the selection threshold is marked as red dashed line. Each dot represents one library variant, and the variants selected for re-screen and sequencing are shown in red.

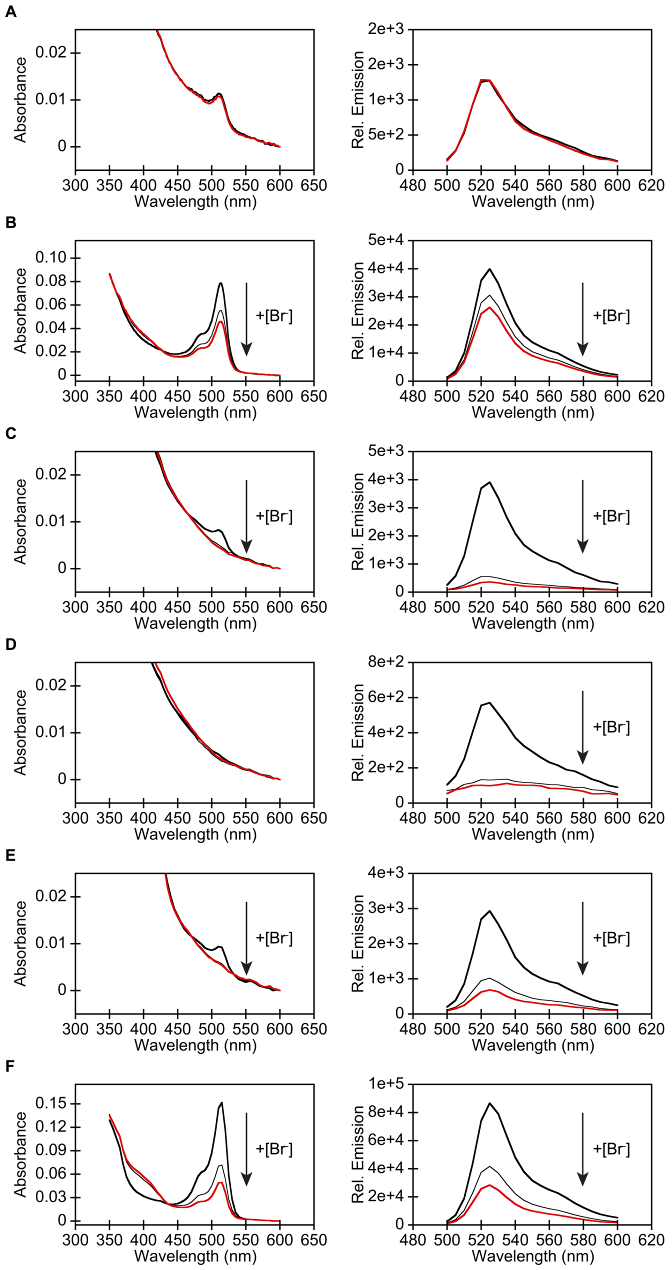


**Figure S10.** Representative absorbance (left) and emission (right) spectra of (A) OFPxm, (B) OFPxm-L69Q-I197T, (C) OFPxm-I68T/L69Q/N144D/I197T (ChlorOFF), (D) OFPxm-V16M K26E/L69Q/A87T/I197T, (E) OFPxm-L69Q/F145L/I197T, and (F) OFPxm-V16A/E34V/L69Q/I197T in *E. coli* lysate suspended in 50 mM sodium phosphate buffer at pH 7 ($\lambda_{abs}$ = 350–600 nm; $\lambda_{ex}$ = 480 nm, $\lambda_{em}$ = 500–600 nm). In panels A and B, the spectra were collected with 0 (bold black), 50, and 100 (red) mM $Br^-$ after the first round of directed evolution. In panels C–F, the spectra were collected with 0 (bold black), 25, and 50 (red) mM $Br^-$ after the second round of directed evolution. Data is shown for one of three biological replicates from the re-screen.

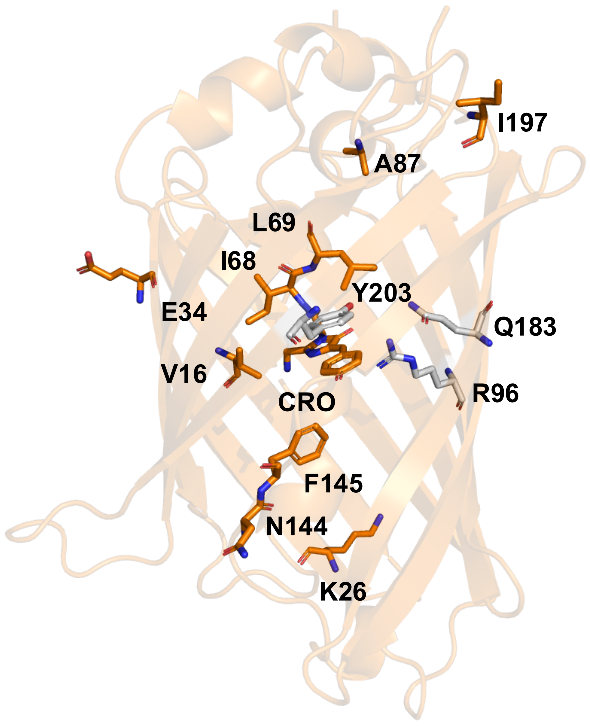


**Figure S11.** Homology model of OFPxm with the chromophore (gray sticks), binding pocket residues (gray sticks), and all mutated residues (orange sticks). Abbreviation: CRO, chromophore.

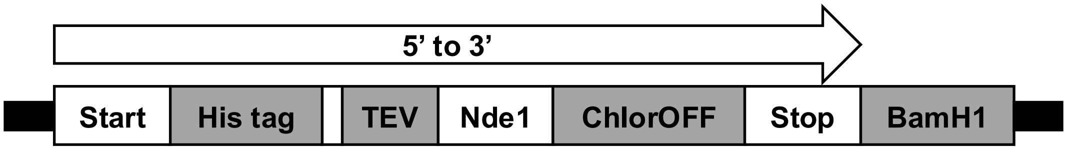


ATGGGCAGCAGCCATCATCATCATCATCACAGCAGCGGCGAGAATCTTTATTTTCAGGGCC
ATATGAGCAAGGGCGAGGAACTGTTCACCGGCGTGGTTCCGATCCTGGTGGAGCTGGAC
GGTGATGTTCACGGCCACAAATTTAGCGTTCGTGGCGAGGGTGAAGGTGATGCGGATTAC
GGCAAGCTGGAAATCAAATTCATTTGCACCACCGGTAAACTGCCGGTTCCGTGGCCGACC
CTGGTTACCACCCTGGGTTACGGCACTCAGTGCTTTGCGCGTTATCCGGAGCACATGAAG
ATGAACGACTTCTTTAAAAGCGCGATGCCGGAGGGTTACATCCAGGAACGTACCATTTTCT
TTCAAGACGATGGCAAGTACAAGACCCGTGGCGAGGTGAAATTCGAAGGTGATACCCTGG
TTAACCGTATCGAGCTGAAGGGCATGGACTTCAAAGAAGATGGTAACATTCTGGGCCACAA
GCTGGAGTACGACTTTAACAGCCACAACGTGTATATCATGCCGGACAAAGCGAACAACGG
TCTGAAGGTTAACTTTAAAATCCGTCACAACATTGAAGGTGGCGGTGTGCAGCTGGCGGAC
CACTACCAAACCAACGTGCCGCTGGGTGATGGCCCGGTTCTGATCCCGACTAACCACTAC
CTGAGCTATCAGACCGCGATTAGCAAGGACCGTAACGAGACCCGTGATCACATGGTGTTC
CTGGAATTCTTTAGCGCGTGCGGTCACACCCACGGCATGGATGAACTGTATAAATAAGGAT
CC

MGSSHHHHHHSSGENLYFQGHMSKGEELFTGVVPILVELDGDVHGHKFSVRGEGEGDADYG
KLEIKFICTTGKLPVPWPTLVTTLGYGTQCFARYPEHMKMNDFFKSAMPEGYIQERTIFFQDDG
KYKTRGEVKFEGDTLVNRIELKGMDFKEDGNILGHKLEYDFNSHNVYIMPDKANNGLKVNFKIR
HNIEGGGVQLADHYQTNVPLGDGPVLIPTNHYLSYQTAISKDRNETRDHMVFLEFFSACGHTH
GMDELYK

**Figure S12.** The pET-28a(+)-TEV-ChlorOFF plasmid design for expression in *E. coli* (top panel), nucleotide sequence (middle panel), and amino acid sequence (bottom panel).

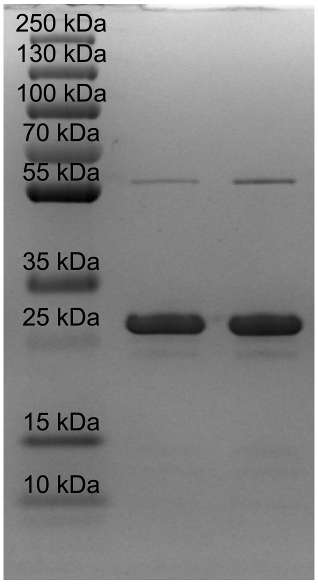


**Figure S13.** (A) A representative Coomassie-stained SDS-PAGE gel of all ChlorOFF protein batches used for characterization in this study. The calculated molecular weight for ChlorOFF is ~29 kDa.

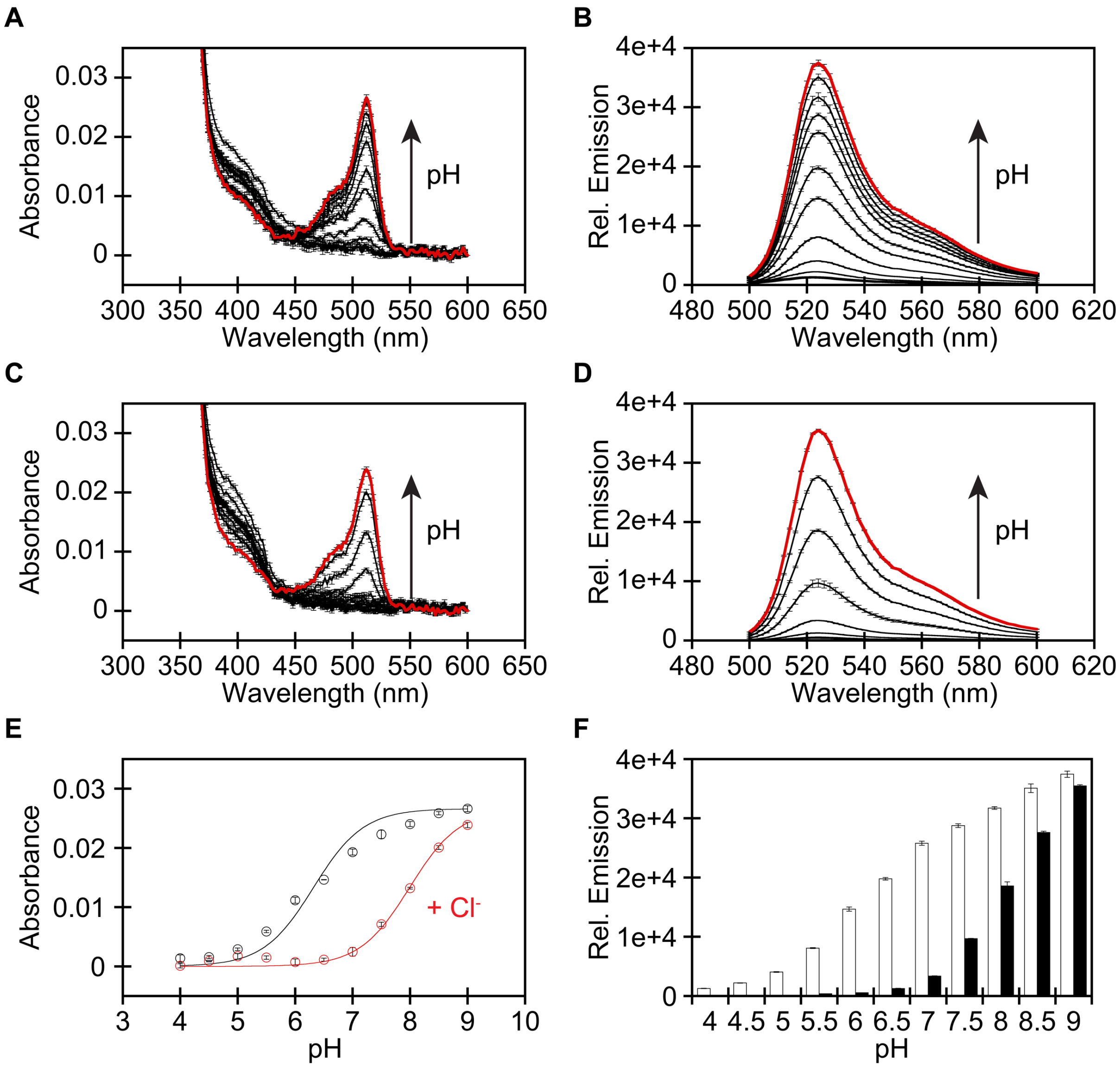


**Figure S14.** (A) Absorbance ($\lambda_{abs}$ = 350–600 nm) and (B) emission ($\lambda_{ex}$ = 480 nm, $\lambda_{em}$ = 500–600 nm) spectra of 3 μM ChlorOFF from pH 4 (bold black) to pH 9 (red). (C) Absorbance and (D) emission spectra of 3 μM ChlorOFF from pH 4 (bold black) to pH 9 (red) in the presence of 100 mM $Cl^-$. (E) Absorbance-dependent p$K_a$ curve of ChlorOFF with 0 mM (black) and 100 mM (red) $Cl^-$ at $\lambda_{abs}$ = 512 nm. The p$K_a$ values in the absence and presence of chloride are 6.32 ± 0.08 and 7.99 ± 0.03, respectively. (F) Emission response of ChlorOFF with 0 mM (white) and 100 mM (black) $Cl^-$ at $\lambda_{em}$ = 524 nm. Data is shown for one of two protein preparations (average with S.E.M.). Data for the second protein preparation is shown in Figure S15. The following buffers were used: 50 mM sodium acetate buffer for pH 4–5.5 or 50 mM sodium phosphate buffer for pH 6–9.

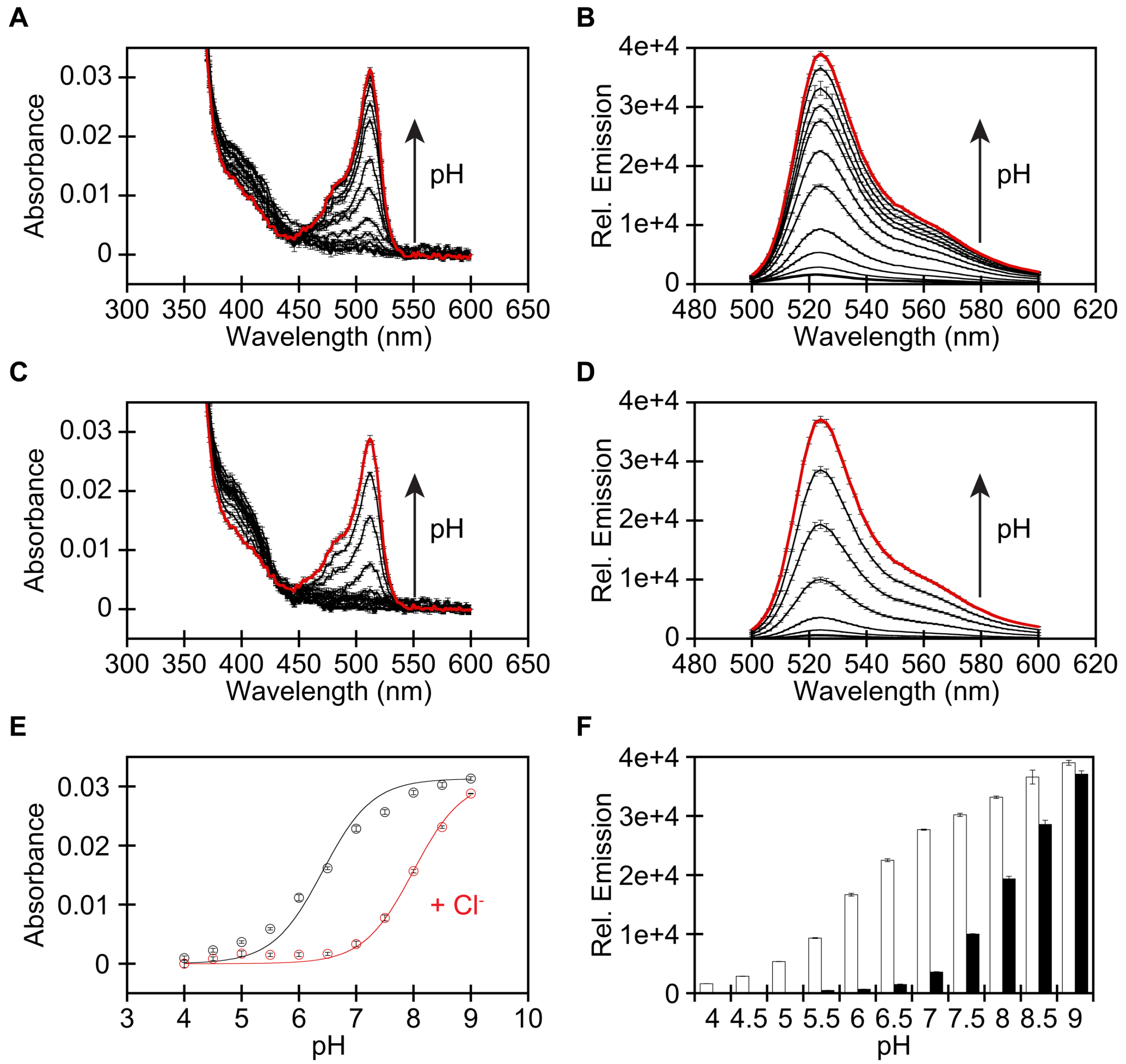


**Figure S15.** (A) Absorbance ($\lambda_{abs}$ = 350–600 nm) and (B) emission ($\lambda_{ex}$ = 480 nm, $\lambda_{em}$ = 500–600 nm) spectra of 3 μM ChlorOFF from pH 4 (bold black) to pH 9 (red). (C) Absorbance and (D) emission spectra of 3 μM ChlorOFF from pH 4 (bold black) to pH 9 (red) in the presence of 100 mM $Cl^-$. (E) Absorbance-dependent p$K_a$ curve of ChlorOFF with 0 mM (black) and 100 mM (red) $Cl^-$ at $\lambda_{abs}$ = 512 nm. The p$K_a$ values in the absence and presence of chloride are 6.42 ± 0.07 and 8.00 ± 0.03, respectively. (F) Emission response of ChlorOFF with 0 mM (white) and 100 mM (black) $Cl^-$ at $\lambda_{em}$ = 524 nm. Data is shown for one of two protein preparations (average with S.E.M.). Data for the second protein preparation is shown in Figure S14. The following buffers were used: 50 mM sodium acetate buffer for pH 4–5.5 or 50 mM sodium phosphate buffer for pH 6–9.

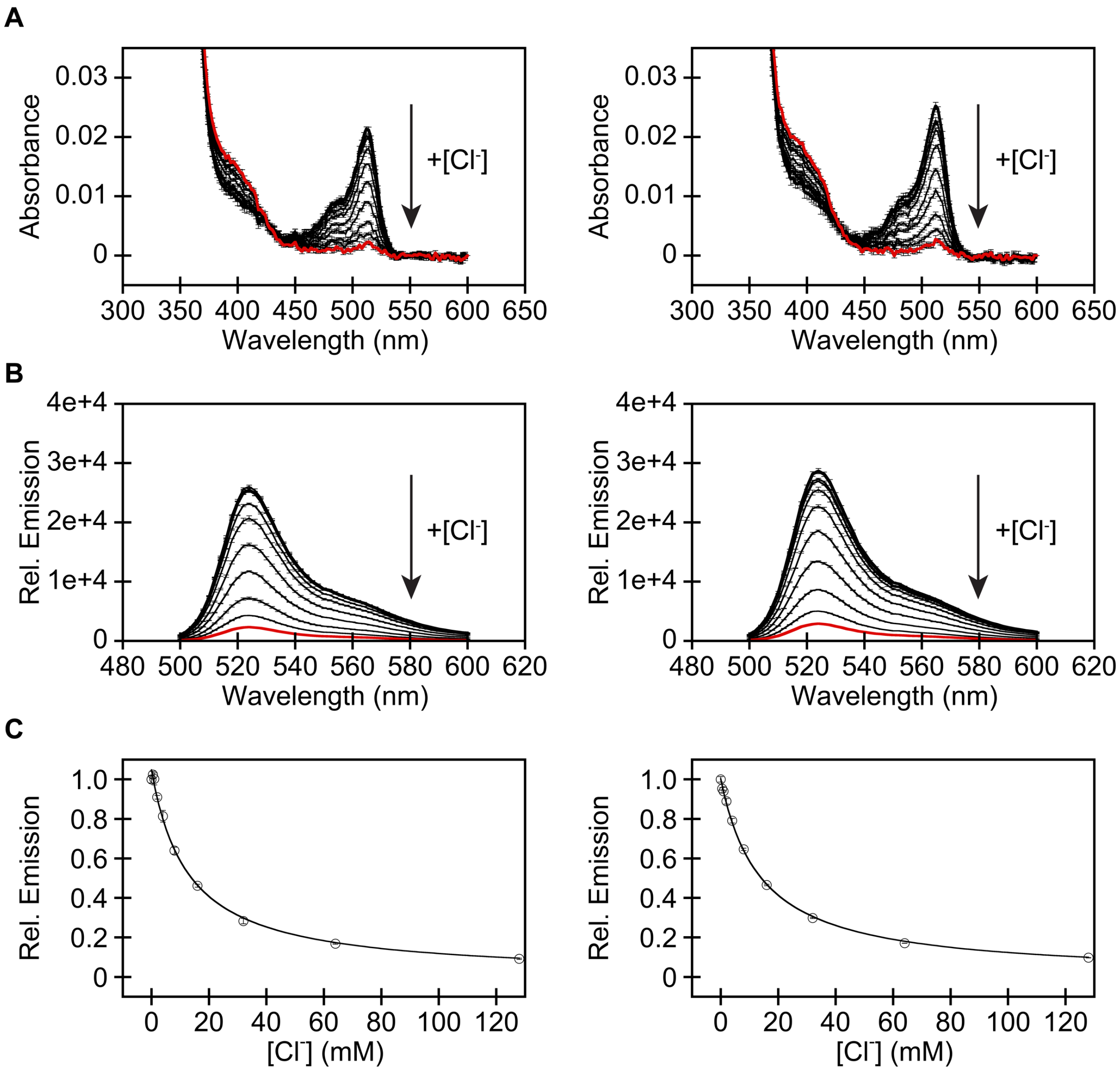


**Figure S16.** (A) Absorbance ($\lambda_{abs}$ = 350–600 nm) and (B) emission ($\lambda_{ex}$ = 480 nm, $\lambda_{em}$ = 500–600 nm) spectra of 3 μM ChlorOFF in 50 mM sodium phosphate buffer at pH 7 in the presence of 0 (bold black), 0.5, 1, 2, 4, 8, 16, 32, 64, and 128 (red) mM $Cl^-$. (C) Emission response ($\lambda_{em}$ = 524 nm) from the titration. The data was normalized to the apo emission and fitted to a single binding site model. For each panel, data is shown for the two protein batches (average with S.E.M.). The apparent dissociation constants ($K_d$) are 12.7 ± 0.7 and 12.7 ± 0.4, respectively.

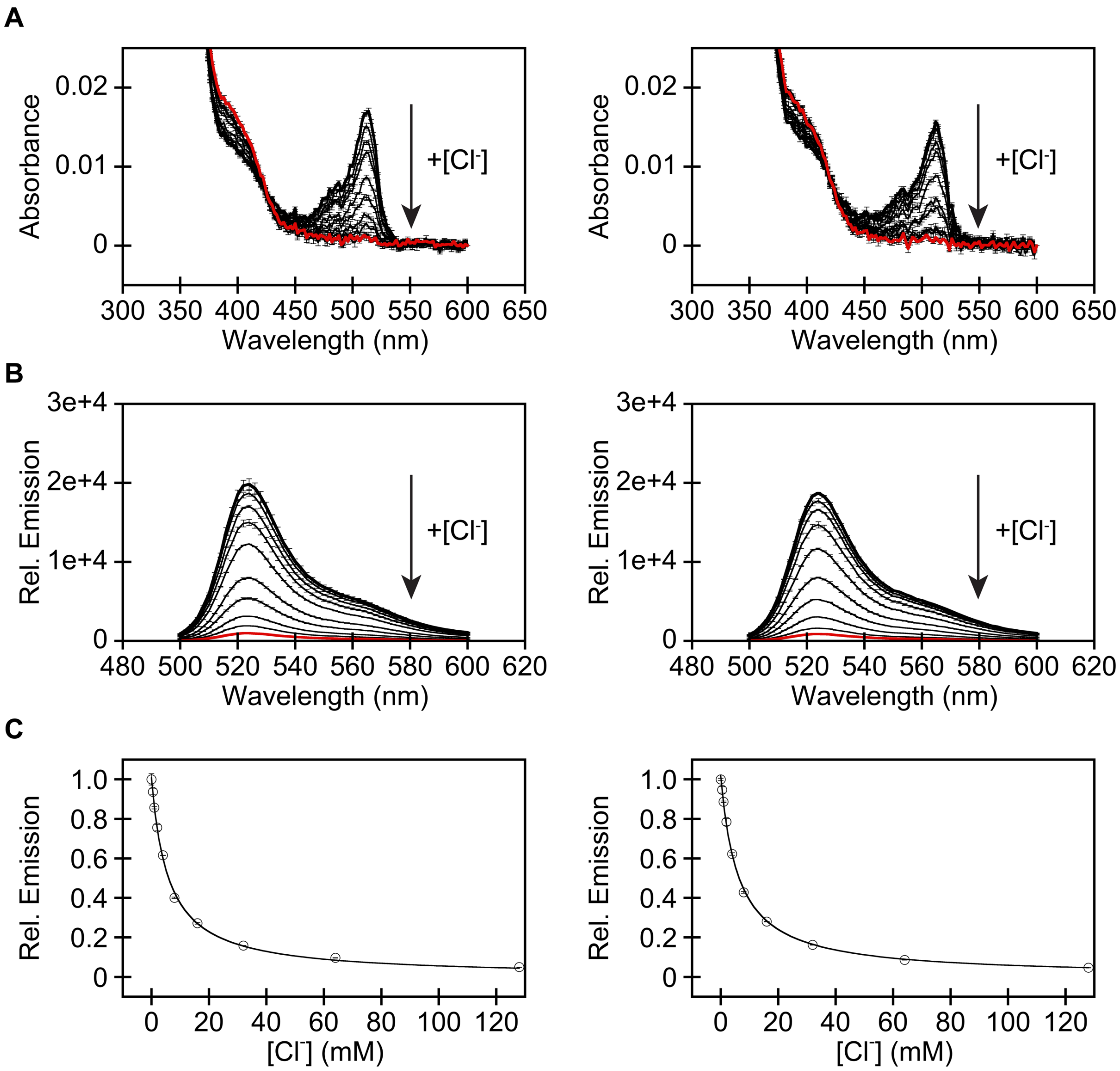


**Figure S17.** (A) Absorbance ($\lambda_{abs}$ = 350–600 nm) and (B) emission ($\lambda_{ex}$ = 480 nm, $\lambda_{em}$ = 500–600 nm) spectra 3 μM ChlorOFF in 50 mM sodium phosphate buffer at pH 6.5 in the presence of 0 (bold black), 0.5, 1, 2, 4, 8, 16, 32, 64, and 128 (red) mM $Cl^-$. (C) Emission response ($\lambda_{em}$ = 524 nm) from the titration. The data was normalized to the apo emission and fitted to a single binding site model. For each panel, data is shown for the two protein preparations (average with S.E.M.). The apparent dissociation constants ($K_d$) are 5.88 ± 0.20 and 6.18 ± 0.19, respectively.

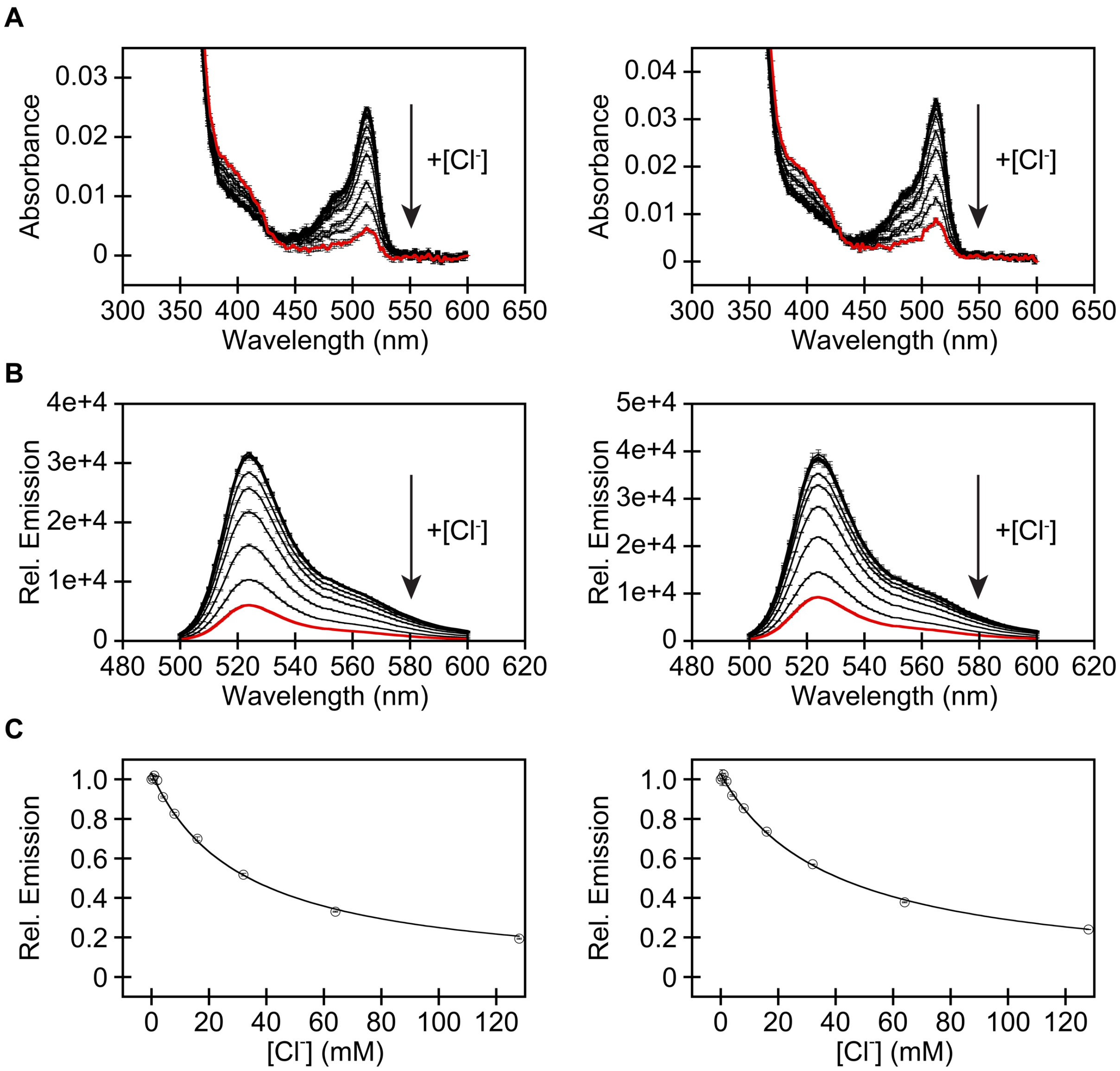


**Figure S18.** (A) Absorbance ($\lambda_{abs}$ = 350–600 nm) and (B) emission ($\lambda_{ex}$ = 480 nm, $\lambda_{em}$ = 500–600 nm) spectra of 3 μM (left) or 3.5 μM (right) ChlorOFF in 50 mM sodium phosphate buffer at pH 7.5 in the presence of 0 (bold black), 0.5, 1, 2, 4, 8, 16, 32, 64, and 128 (red) mM $Cl^-$. (C) Emission response ($\lambda_{em}$ = 524 nm) from the titration. The data was normalized to the apo emission and fitted to a single binding site model. For each panel, data is shown for the two protein batches (average with S.E.M.). The apparent dissociation constants ($K_d$) are 31.95 ± 1.52 and 31.32 ± 2.32, respectively.

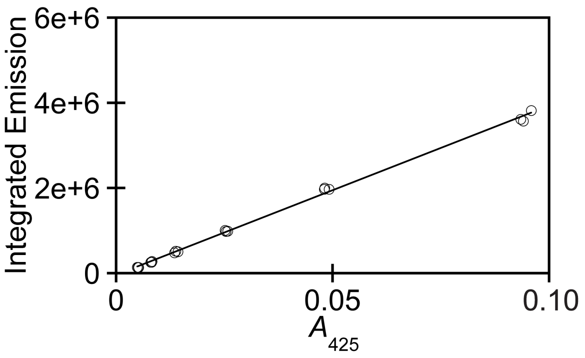


**Figure S19.** Plot of the integrated emission ($\lambda_{ex}$ = 425 nm, $\lambda_{em}$ = 440–800 nm) versus the corrected absorbance intensity at 425 nm for Coumarin 135 in 50% ethanol ($R^2$ > 0.95) to determine the fluorescence quantum yield. Data for each concentration was measured in triplicate.

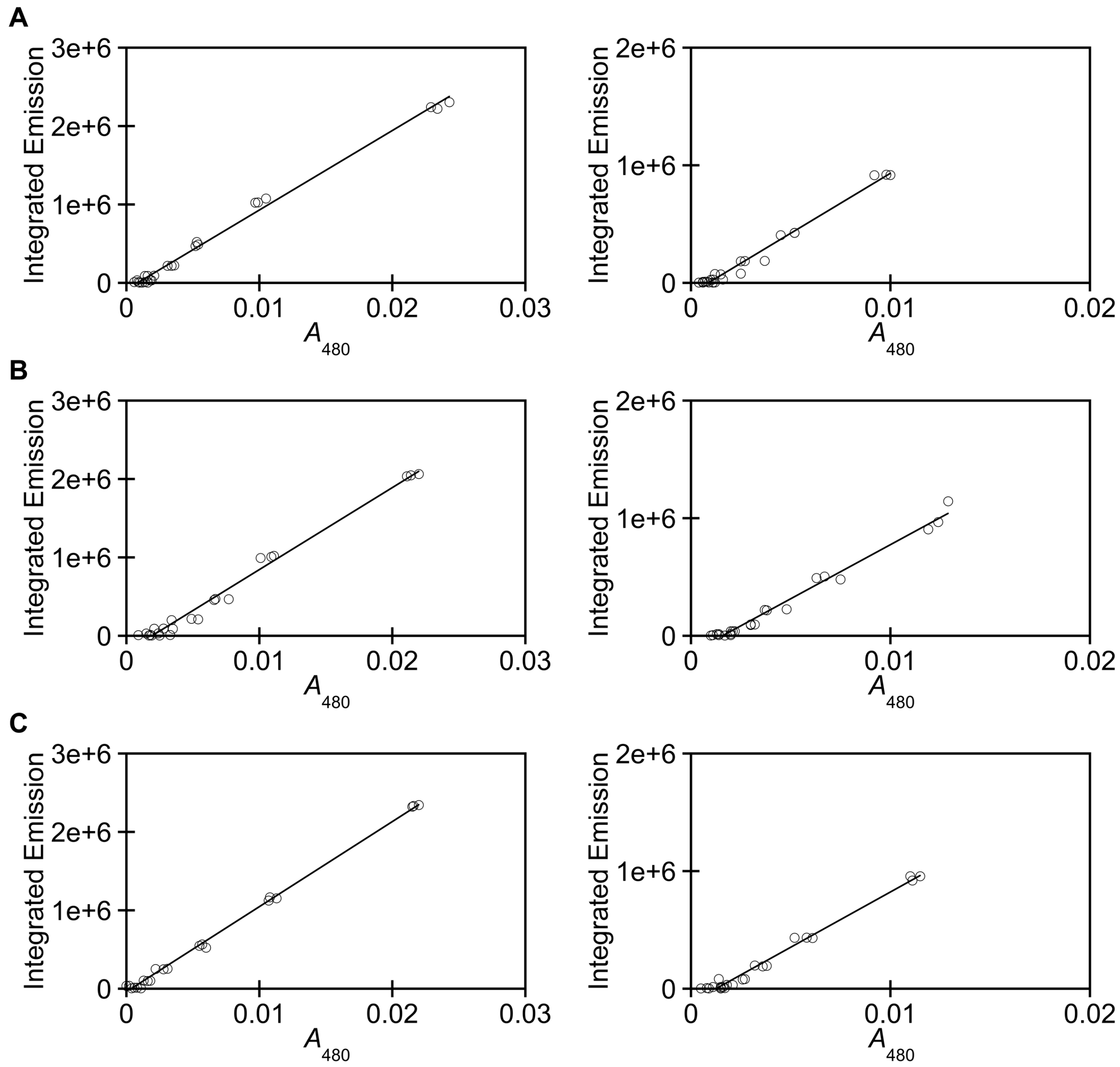


**Figure S20.** Plot of the integrated emission ($\lambda_{ex}$ = 480 nm, $\lambda_{em}$ = 500–600 nm) versus the corrected absorbance intensity at 480 nm for ChlorOFF in 50 mM sodium phosphate buffer at (A) pH 6.5, (B) pH 7, and (C) pH 7.5 to determine the fluorescence quantum yield ($R^2$ > 0.95). Data for each concentration was measured in triplicate. For each panel, all data is shown for the two protein batches.

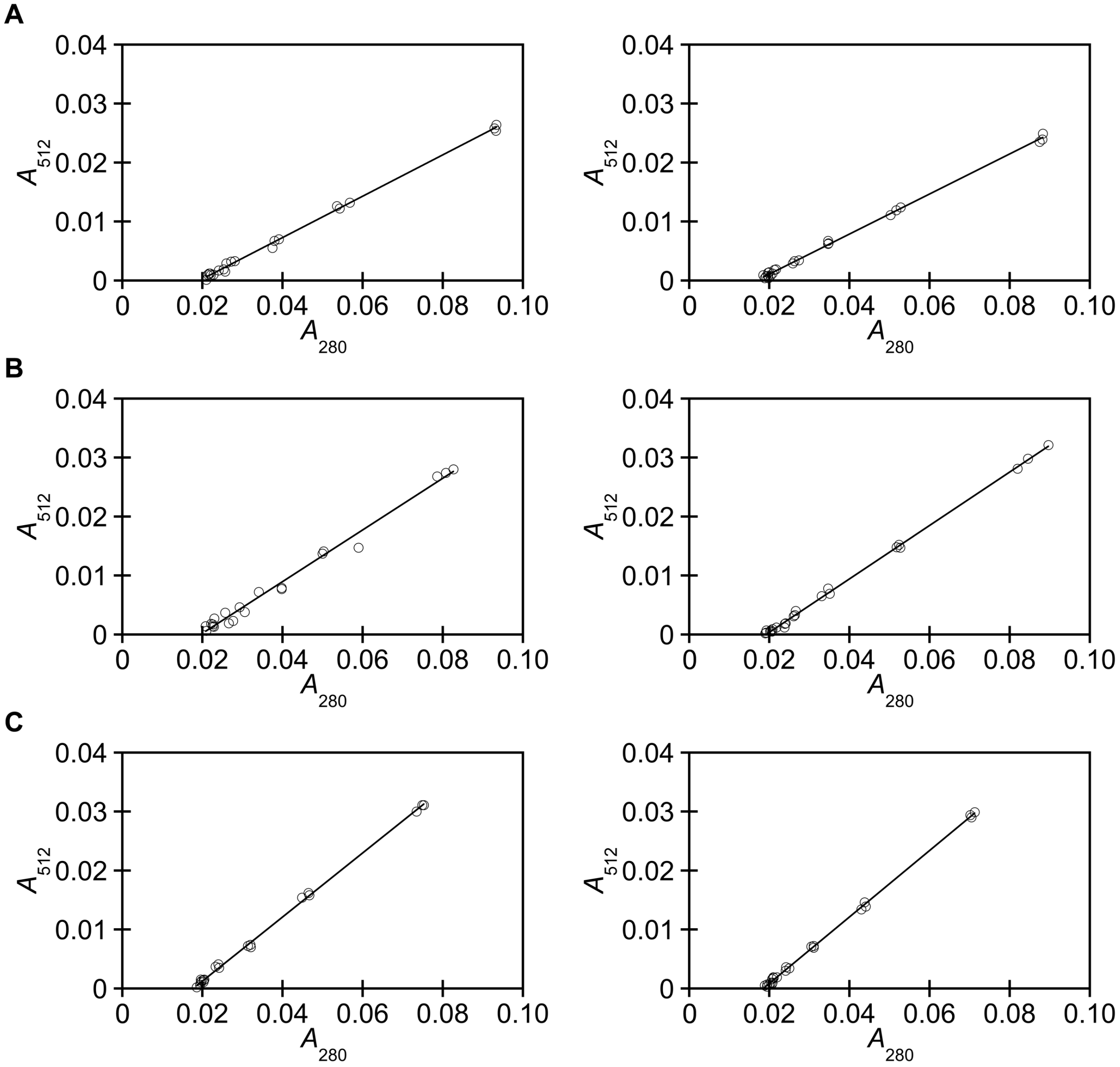


**Figure S21.** The corrected absorbance intensity at 512 nm is plotted versus the corrected absorbance intensity at 280 nm for ChlorOFF in 50 mM sodium phosphate buffer at (A) pH 6.5, (B) pH 7, and (C) pH 7.5 to determine the extinction coefficient ($R^2$ > 0.99). Data for each concentration was measured in triplicate. For each panel, all data is shown for the two protein batches.

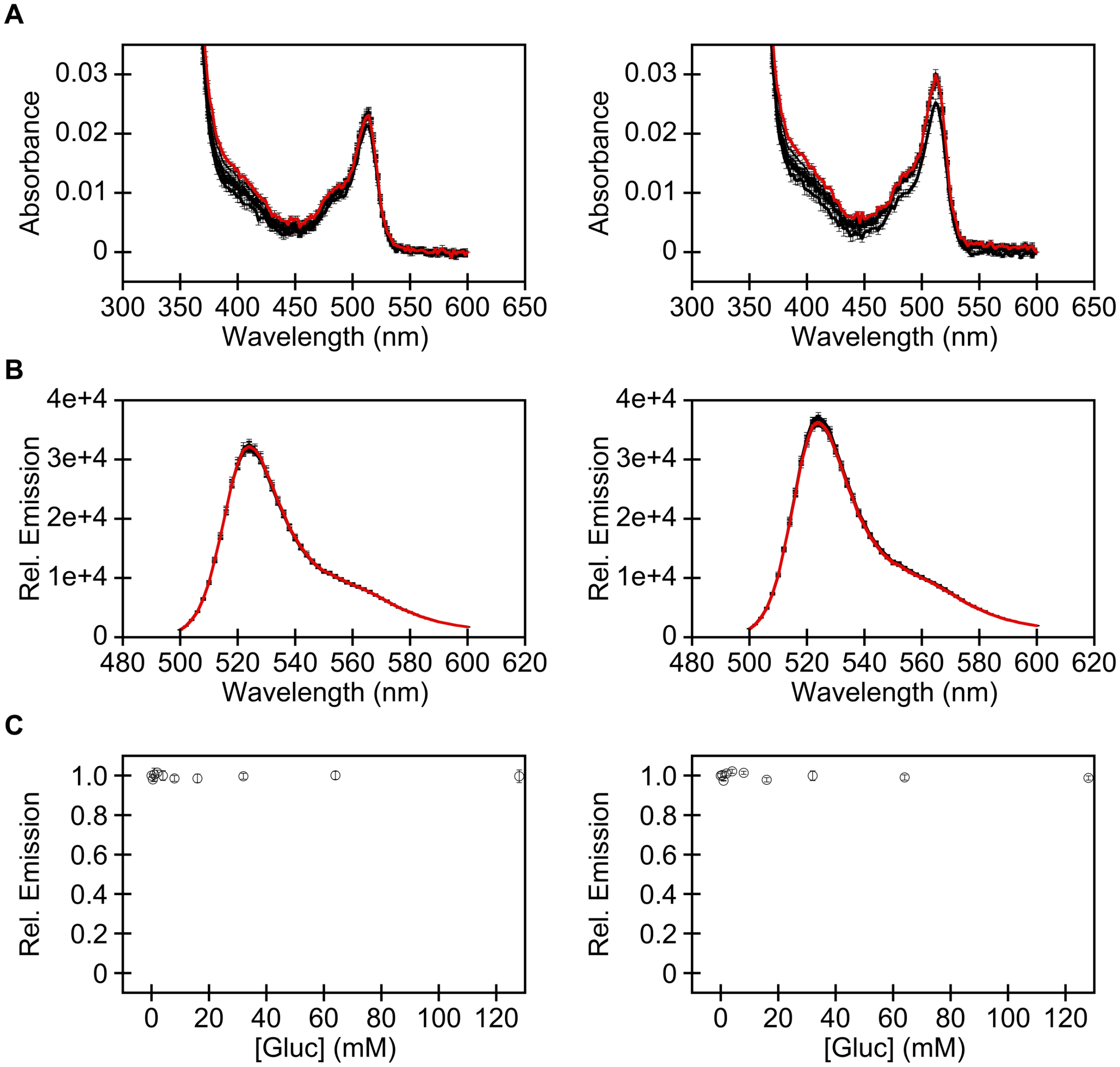


**Figure S22.** (A) Absorbance ($\lambda_{abs}$ = 350–600 nm) and (B) emission ($\lambda_{ex}$ = 480 nm, $\lambda_{em}$ = 500–600 nm) spectra of 3 µM (left) or 3.5 µM (right) ChlorOFF in 50 mM sodium phosphate buffer at pH 7 in the presence of 0 (bold black), 0.5, 1, 2, 4, 8, 16, 32, 64, and 128 (red) mM gluconate (Gluc). (C) Emission response ($\lambda_{em}$ = 524 nm) from the titration. The data was normalized to the apo emission. For each panel, data is shown for the two protein batches (average with S.E.M.). The apparent dissociation constants ($K_d$) could not be determined.

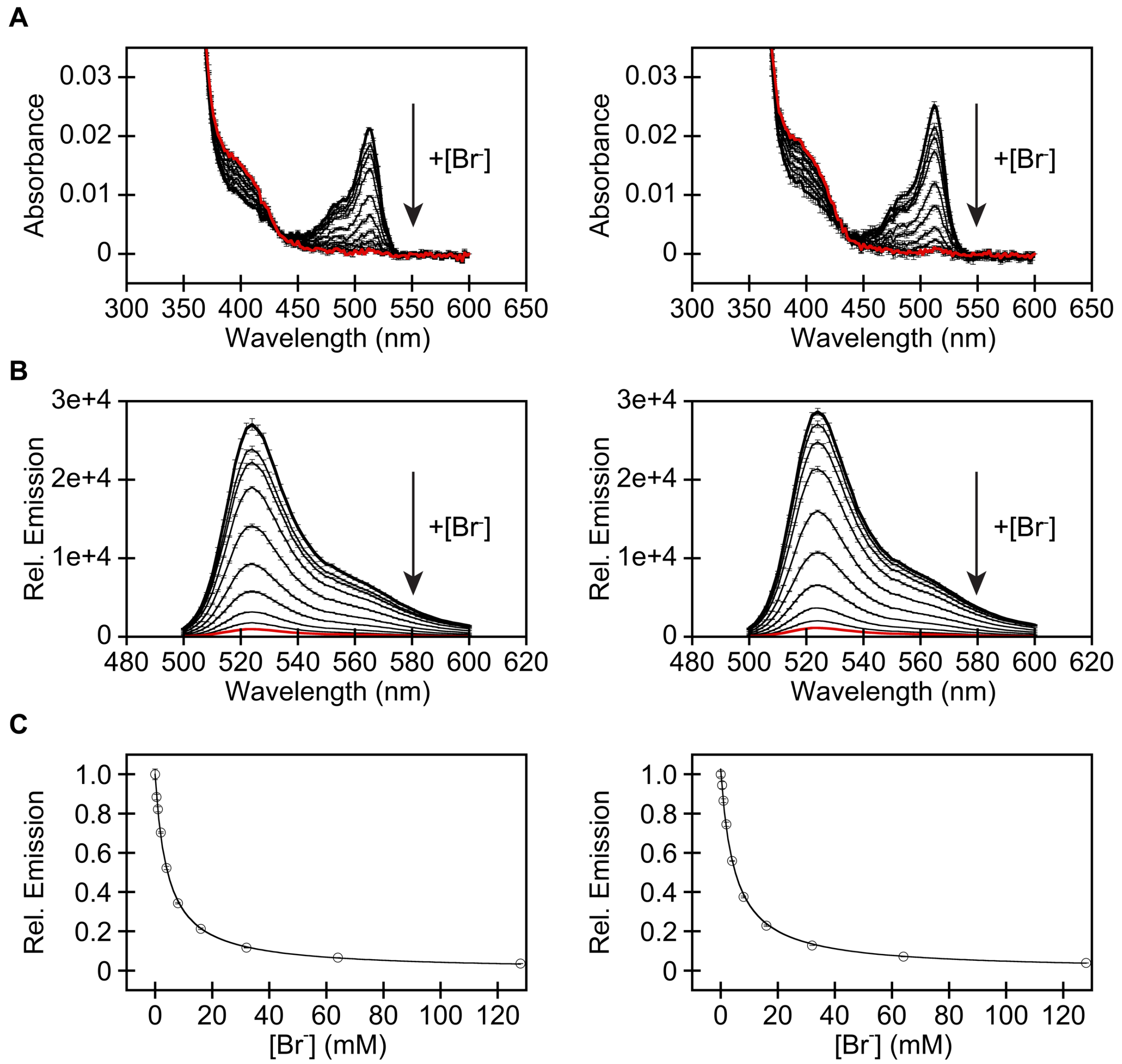


**Figure S23.** (A) Absorbance ($\lambda_{abs}$ = 350–600 nm) and (B) emission ($\lambda_{ex}$ = 480 nm, $\lambda_{em}$ = 500–600 nm) spectra of 3 μM ChlorOFF in 50 mM sodium phosphate buffer at pH 7 in the presence of 0 (bold black), 0.5, 1, 2, 4, 8, 16, 32, 64, and 128 (red) mM $Br^-$. (C) Emission response ($\lambda_{em}$ = 524 nm) from the titration. The data was normalized to the apo emission and fitted to a single binding site model. For each panel, data is shown for the two protein batches (average with S.E.M.). The apparent dissociation constants ($K_d$) are 4.39 ± 0.11 and 4.86 ± 0.20, respectively.

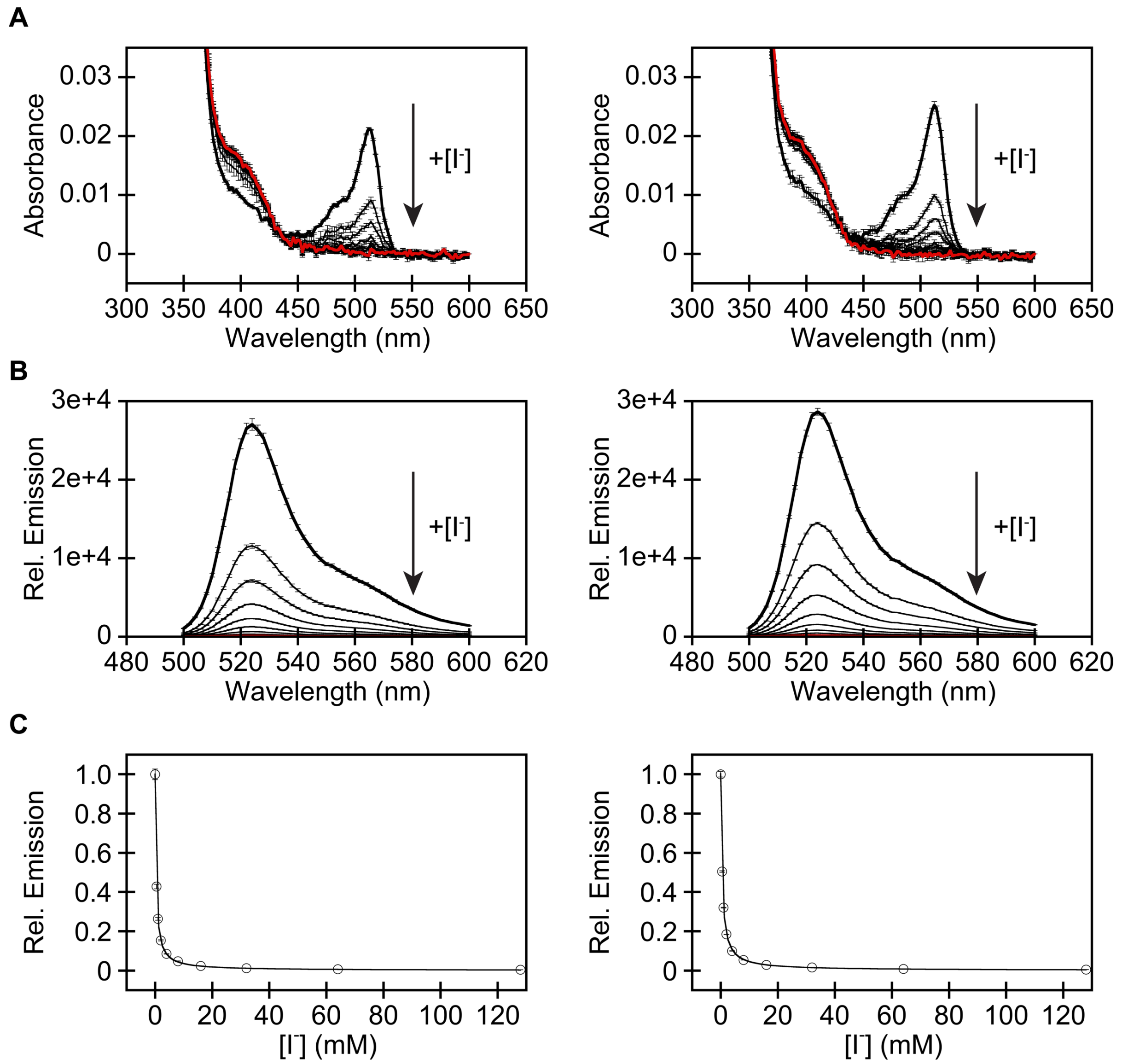


**Figure S24.** (A) Absorbance ($\lambda_{abs}$ = 350–600 nm) and (B) emission ($\lambda_{ex}$ = 480 nm, $\lambda_{em}$ = 500–600 nm) of 3 μM ChlorOFF in 50 mM sodium phosphate buffer pH 7 in the presence of 0 (bold black), 0.5, 1, 2, 4, 8, 16, 32, 64, and 128 (red) mM $I^-$. (C) Emission response ($\lambda_{em}$ = 524 nm) from the titration. The data was normalized to the apo emission and fitted to a single binding site model. For each panel, data is shown for the two protein batches (average with S.E.M.). The apparent dissociation constants ($K_d$) are 0.37 ± 0.003 and 0.48 ± 0.01, respectively.

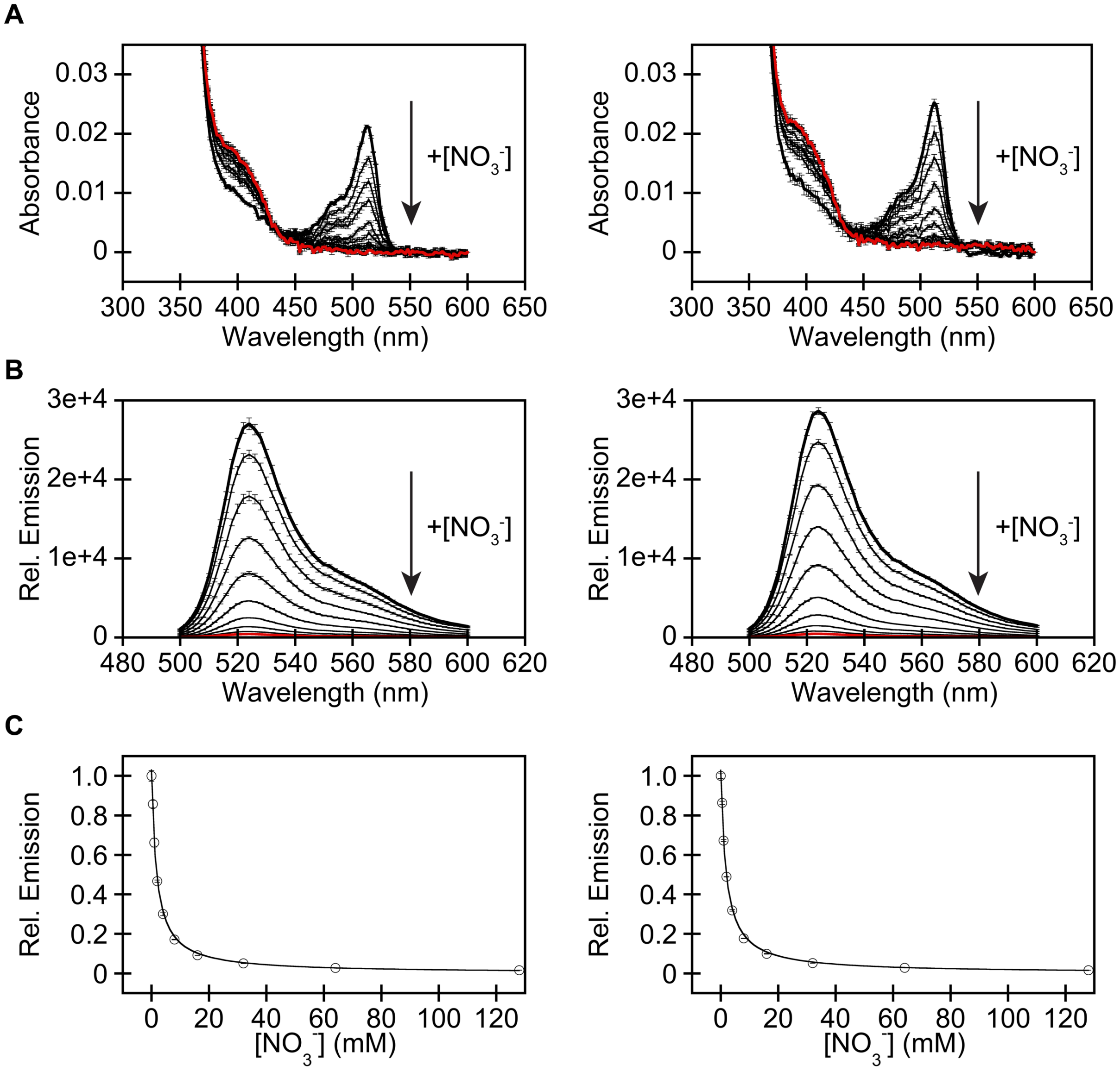


**Figure S25.** (A) Absorbance ($\lambda_{abs}$ = 350–600 nm) and (B) emission ($\lambda_{ex}$ = 480 nm, $\lambda_{em}$ = 500–600 nm) spectra of 3 µM ChlorOFF in 50 mM sodium phosphate buffer at pH 7 in the presence of 0 (bold black), 0.5, 1, 2, 4, 8, 16, 32, 64, and 128 (red) mM $NO_3^-$. (C) Emission response ($\lambda_{em}$ = 524 nm) from the titration. The data was normalized to the apo emission and fitted to a single binding site model. For each panel, data is shown for the two protein batches (average with S.E.M.). The apparent dissociation constants ($K_d$) are 1.77 ± 0.12 and 1.88 ± 0.12, respectively.

**Table S4.** *In vitro* spectroscopic properties of ChlorOFF. Data is shown as the average of the two protein batches with S.E.M. Abbreviation: N.D., not determined.

| | | pH 6.5 | pH 7 | pH 7.5 |
|---|---|---|---|---|
| **Molar Extinction coefficient ($M^{-1}cm^{-1}$)** | **Apo** | 8,108 ± 130 | 10,476 ± 176 | 13,005 ± 237 |
| **Quantum yield** | **Apo** | 0.61 ± 0.001 | 0.59 ± 0.04 | 0.61 ± 0.04 |
| **Molar brightness** | **Apo** | 4.96 ± 0.06 | 6.19 ± 0.29 | 7.91 ± 0.42 |
| **$K_d$ (mM)** | **$Cl^-$** | 6.03 ± 0.10 | 12.7 ± 0.2 | 31.6 ± 0.7 |
| | **$Br^-$** | N.D. | 4.63 ± 0.12 | N.D. |
| | **$I^-$** | N.D. | 0.43 ± 0.02 | N.D. |
| | **$NO_3^-$** | N.D. | 1.83 ± 0.05 | N.D. |

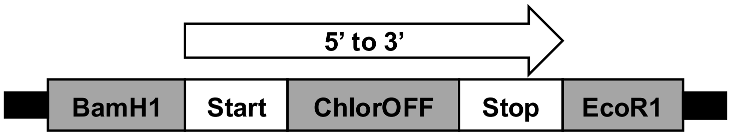


GGATCCATGTCTAAGGGGGAGGAGCTGTTTACTGGCGTGGTTCCGATCCTGGTGGAGCTG
GACGGTGATGTTCACGGCCACAAATTTAGCGTTCGTGGCGAGGGTGAAGGTGATGCGGAT
TACGGCAAGCTGGAAATCAAATTCATTTGCACCACCGGTAAACTGCCGGTTCCGTGGCCG
ACCCTGGTTACCACCCTGGGTTACGGCATTCTGTGCTTTGCGCGTTATCCGGAGCACATGA
AGATGAACGACTTCTTTAAAAGCGCGATGCCGGAGGGTTACATCCAGGAACGTACCATTTT
CTTTCAAGACGATGGCAAGTACAAGACCCGTGGCGAGGTGAAATTCGAAGGTGATACCCT
GGTTAACCGTATCGAGCTGAAGGGCATGGACTTCAAAGAAGATGGTAACATTCTGGGCCA
CAAGCTGGAGTACAACTTTAACAGCCACAACGTGTATATCATGCCGGACAAAGCGAACAAC
GGTCTGAAGGTTAACTTTAAAATCCGTCACAACATTGAAGGTGGCGGTGTGCAGCTGGCG
GACCACTACCAAACCAACGTGCCGCTGGGTGATGGCCCGGTTCTGATCCCGATTAACCAC
TACCTGAGCTATCAGACCGCGATTAGCAAGGACCGTAACGAGACCCGTGATCACATGGTG
TTCCTGGAATTCTTTAGCGCGTGCGGGCACACCCACGGAATGGACGAGCTGTATAAGTGA
GAATTC

MSKGEELFTGVVPILVELDGDVHGHKFSVRGEGEGDADYGKLEIKFICTTGKLPVPWPTLVTTL
GYGILCFARYPEHMKMNDFFKSAMPEGYIQERTIFFQDDGKYKTRGEVKFEGDTLVNRIELKG
MDFKEDGNILGHKLEYNFNSHNVYIMPDKANNGLKVNFKIRHNIEGGGVQLADHYQTNVPLGD
GPVLIPINHYLSYQTAISKDRNETRDHMVFLEFFSACGHTHGMDELYK

**Figure S26.** The pcDNA3.1(+)-ChlorOFF plasmid design for expression in mammalian cells (top panel), nucleotide sequence (middle panel), and amino acid sequence (bottom panel).

```
run("Concatenate...", "all_open open");
run("Split Channels");
selectWindow("green");
run("StackReg", "transformation=Translation");//align images
run("Despeckle");
run("Remove Outliers...", "radius=3 threshold=500 which=Bright");//reduce intensity from debris
run("Subtract Background...", "rolling=75 stack");
run("Merge Channels...", "c2=green c4=dic create");//merge channels as single stack
Property.set("CompositeProjection", "null");
Stack.setDisplayMode("color");

run("Split Channels");
selectWindow("green");
run("Z Project...", "projection=[Max Intensity]");
run("Threshold...");
setThreshold(100, 6000);//Default threshold
waitForUser("Apply Threshold");
run("Convert to Mask");//create mask
run("Watershed");//delineate between cells and cell clusters
run("Analyze Particles...", "size=10-Infinity exclude clear add");//create ROIs
waitForUser("Check ROIs");//remove ROIs with floating debris or moving cells
roiManager("multi measure");//apply ROIs to Green channel and measure Median Intensity
```

**Figure S27.** ImageJ script used to analyze the fluorescence microscopy images of U-2 OS cells expressing ChlorOFF.

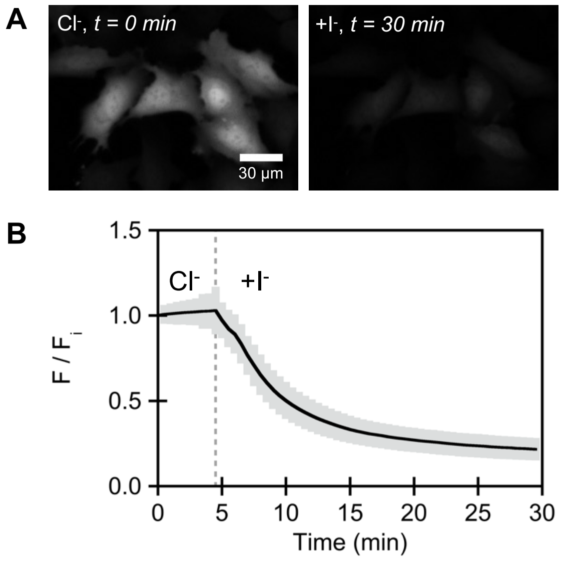


**Figure S28.** Characterization of ChlorOFF in the U-2 OS cell model with widefield fluorescence microscopy. (a) Representative fluorescence images for the exchange assay with chloride-iodide (*n* = 1422). (b) The average median fluorescence intensity with standard deviation is reported for the *n* regions of interest from three biological replicates. The vertical dashed line is the start of the perfusion for the anion exchange. All experiments were conducted at 37 °C in a PBS buffer at pH 7.4 (see *Methods*) containing 137 mM NaCl or 68.5 mM NaCl/68.5 mM NaI. Scale bar = 30 μm.

```
run("Concatenate...", "all_open open");
run("Split Channels");
selectWindow("Cyan");
run("StackReg", "transformation=Translation");//align images
run("Despeckle");
run("Remove Outliers...", "radius=3 threshold=500 which=Bright");//reduce intensity from debris
run("Subtract Background...", "rolling=75 stack");
selectWindow("Red");
run("StackReg", "transformation=Translation");//align images
run("Despeckle");
run("Remove Outliers...", "radius=3 threshold=500 which=Bright");//reduce intensity from debris
run("Subtract Background...", "rolling=75 stack");
run("Merge Channels...", "c1=Red c4=DIC c5=Cyan create");//merge channels as single stack
Property.set("CompositeProjection", "null");
Stack.setDisplayMode("color");

run("Duplicate...", "title=stack duplicate frames=1-27");//all frames except pH 8 clamping slice
run("Split Channels");
selectWindow("Cyan");
run("Z Project...", "projection=[Max Intensity]");
run("Threshold...");
setThreshold(750, 6000);//Default threshold
waitForUser("Apply Threshold");
setOption("BlackBackground", false);
run("Convert to Mask");//create mask
run("Watershed");//delineate between cells and cell clusters
run("Analyze Particles...", "size=40-Infinity exclude clear include add");//create ROIs
waitForUser("Check ROIs");//remove ROIs with floating debris or moving cells
roiManager("multi measure");//apply ROIs to Red and Cyan channels with all frames to measure
Median Intensity
```

**Figure S29.** ImageJ script used to analyze the fluorescence microscopy images of U-2 OS cells stained with BCECF.

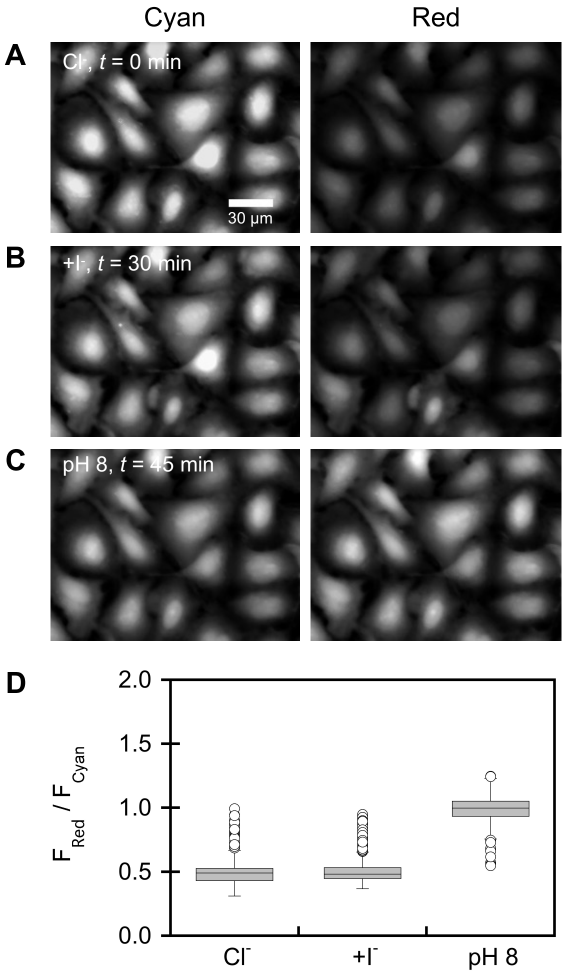


**Figure S30.** Characterization of BCECF in the U-2 OS cell model with widefield fluorescence microscopy. Representative fluorescence images from the exchange assay with chloride-iodide at (a) $t$ = 0 min and (b) $t$ = 30 min ($\Delta$F = 1.07 $\pm$ 0.2) and from the pH 8 ionophore clamp at (c) $t$ = 45 min ($\Delta$F = 2.09 $\pm$ 0.3, $n$ = 2265). Images are shown for the BCECF emission response with excitation at 436 nm (Cyan) and at 495 nm (Red). (d) For each condition, the average median fluorescence intensity with standard deviation is reported for the $n$ regions of interest from three biological replicates. All experiments were conducted at 37 °C in a PBS buffer (see *Methods*). Scale bar = 30 µm.

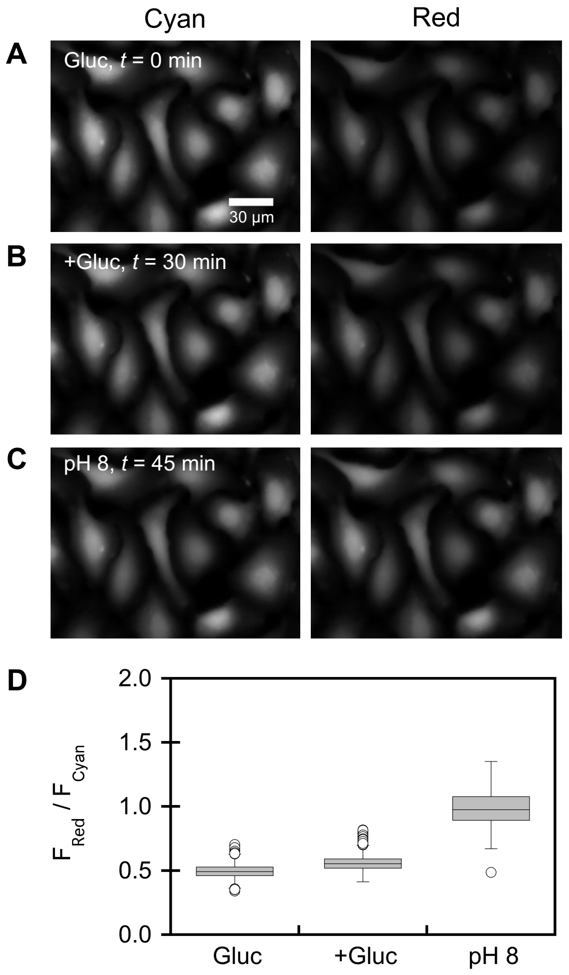


**Figure S31.** Characterization of BCECF in the U-2 OS cell model with widefield fluorescence microscopy. Representative fluorescence images from the exchange assay with gluconate-gluconate at (a) $t$ = 0 min and (b) $t$ = 30 min ($\Delta F = 1.13 \pm 0.1$) and from the pH 8 ionophore clamp at (c) $t$ = 45 min ($\Delta F = 1.99 \pm 0.2$, $n$ = 2247). Images are shown for the BCECF emission response with excitation at 436 nm (Cyan) and at 495 nm (Red). (d) For each condition, the average median fluorescence intensity with standard deviation is reported for the $n$ regions of interest from three biological replicates. All experiments were conducted at 37 °C in a PBS buffer (see *Methods*). Scale bar = 30 µm. Abbreviation: Gluc, gluconate.

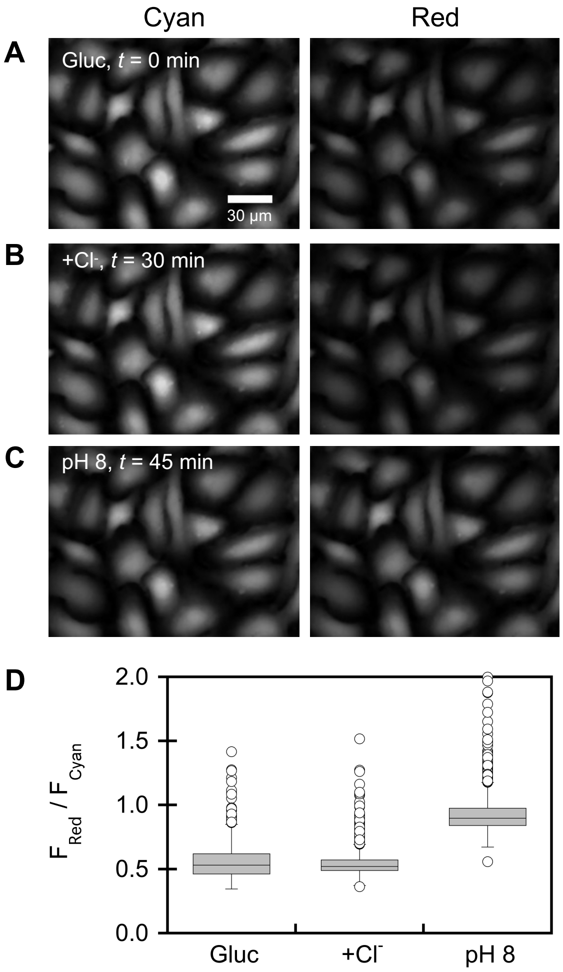


**Figure S32.** Characterization of BCECF in the U-2 OS cell model with widefield fluorescence microscopy. Representative fluorescence images from the exchange assay with gluconate-chloride at (a) $t$ = 0 min and (b) $t$ = 30 min ($\Delta$F = 1.01 $\pm$ 0.1) and from the pH 8 ionophore clamp at (c) $t$ = 45 min ($\Delta$F = 1.70 $\pm$ 0.2, $n$ = 2467). Images are shown for the BCECF emission response with excitation at 436 nm (Cyan) and at 495 nm (Red). (d) For each condition, the average median fluorescence intensity with standard deviation is reported for the $n$ regions of interest from three biological replicates. All experiments were conducted at 37 °C with a PBS buffer (see *Methods*). Scale bar = 30 µm. Abbreviation: Gluc, gluconate.

**Table S5.** Summary of mutations in fluorescent protein indicators for chloride.

| Entry | Name of Indicator | Parent | Mutations Relative to Parent | Main Text Reference |
|---|---|---|---|---|
| 1 | YFP | GFP from jellyfish *Aequorea victoria* | S65G/V68L/S72A/T203Y | 7 |
| 2 | YFP-H148Q | YFP | H148Q | 7 |
| 3 | YFP-H148Q-V150T | YFP-H148Q | V150T | 14 |
| 4 | YFP-H148Q-I152L | YFP-H148Q | I152L | 14 |
| 5 | YFP-H148Q-V163S | YFP-H148Q | V163S | 14 |
| 6 | ClsM | YFP | F46L/Q69K/H148Q/I152L/V163S | 13 |
| 7 | Cl-YFP | ClsM | S175G/S205V | 13 |
| 8 | mCl-YFP | Cl-YFP | A206K | 13 |
| 9 | Cl-sensor | YFP | H148Q/I152L/V163S | 12 |
| 10 | Clomeleon chloride-sensitive YFP Topaz domain (CT) | avGFP | S65G/S72A/K79R/T203Y/H231L | 11 |
| 11 | CT-Q69T | CT | Q69T | 10 |
| 12 | CT-H148Q | CT | H148Q | 10 |
| 13 | CT-Q183A | CT | Q183A | 10 |
| 14 | CT-Q69T-V163A | CT | Q69T/V163A | 10 |
| 15 | CT-H148Q-V163S | CT | H148Q/V163S | 10 |
| 16 | CT-V150A-V163A | CT | V150A/V163A | 10 |
| 17 | CT-H148Q-I152L-V163S | CT | H148Q/I152L/V163S | 10 |
| 18 | CT-H148Q-V163A-L201I | CT | H148Q/V163A/L201I | 10 |
| 19 | E1GFP | EGFP | S65T | 15 |
| 20 | E2GFP | EGFP | S65T/T203Y | 15 |
| 21 | ClopHensor-H148G-V224L | ClopHensor chloride-sensitive E2GFP domain | H148G/V224L | 16 |
| 22 | LSSmsfClopHensor | Superfolder GFP | T203Y | 9 |
| 23 | mNeonGreen-R195Y | mNeonGreen | R195Y | 23 |
| 24 | ChlorON-1 | mNeonGreen | K143W/R195L | 24 |
| 25 | ChlorON-2 | mNeonGreen | K143R/R195I | 24 |
| 26 | ChlorON-3 | mNeonGreen | K143R/R195L | 24 |
| 27 | mBeRFP-D162S | mBeRFP | D162S | 20 |
| 28 | mBeRFP-D162A | mBeRFP | D162A | 20 |
| 29 | mBeRFP-S94V | mBeRFP | S94V | 20 |
| 30 | mBeRFP-S94V | mBeRFP | R205Y | 20 |
| 31 | mBeRFP-R205Y | mBeRFP | D162S/S94V | 20 |
| 32 | mBeRFP-S94V/R205Y | mBeRFP | S94V/R205Y | 20 |
| 33 | mBeRFP-D162S/R205Y | mBeRFP | D162S/R205Y | 20 |
| 34 | mBeRFP-D162S-S94V-R205Y | mBeRFP | D162S/S94V/R205Y | 20 |
| 35 | OFPxm-L69Q/I197T | OFPxm | L69Q/I197T | *This study* |
| 36 | OFPxm-V16A-E34A | OFPxm-L69Q/I197T | V16A/E34A | *This study* |
| 37 | OFPxm-F145L | OFPxm-L69Q/I197T | F145L | *This study* |
| 38 | OFPxm-V16M-K26E-A87T | OFPxm-L69Q/I197T | V16M/K26E/A87T | *This study* |
| 39 | ChlorOFF | OFPxm-L69Q/I197T | I68T/N144D | *This study* |

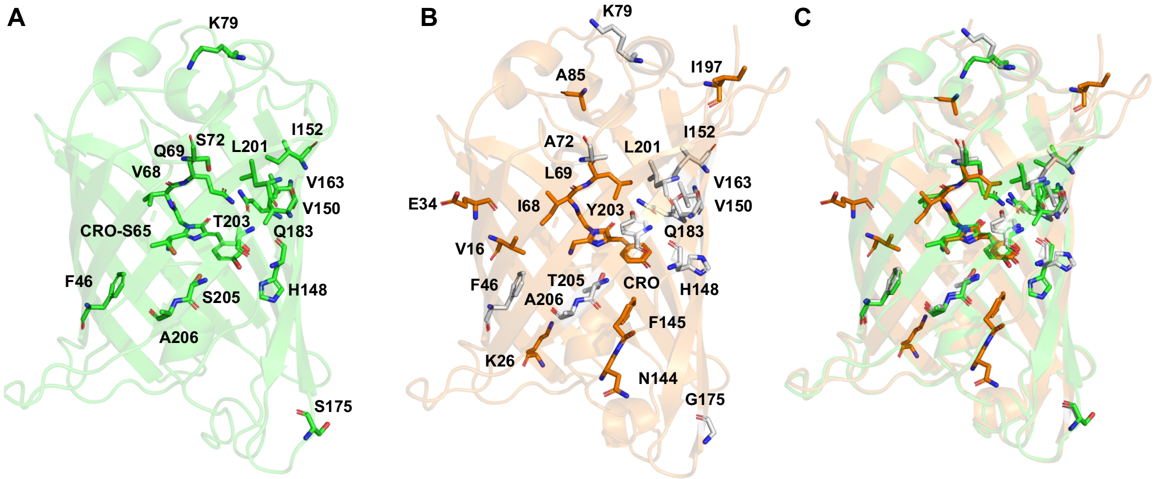


**Figure S33.** (A) Structure of GFP (PDB ID: 1EMA) with the chromophore (green sticks) and mutated residues (green sticks) in Table S5 for entries 1–18. Note: H231 is not present in the structure. (B) Homology model of OFPxm with the chromophore (orange sticks), all mutated residues (orange sticks), homologous mutated residues from panel A (gray sticks). Note: The homologous H231 residue is not shown. (C) Overlay of panels A and B. Abbreviation: CRO, chromophore.

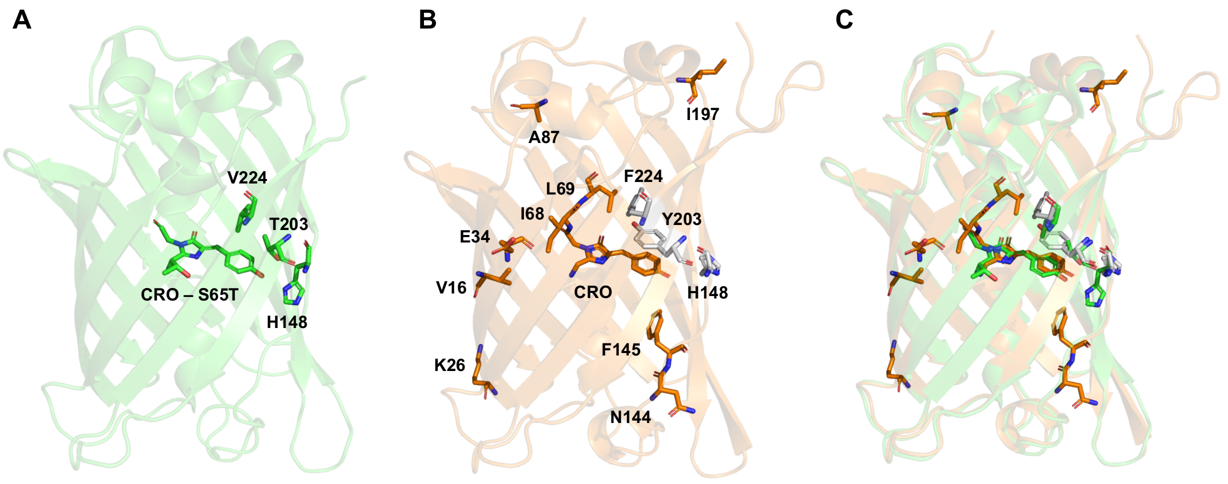


**Figure S34.** (A) Structure of EGFP (PDB ID: 2Y0G) with the chromophore (green sticks) and mutated residues (green sticks) in Table S5 for entries 19–21. (B) Homology model of OFPxm with the chromophore (orange sticks), all mutated residues (orange sticks), homologous mutated residues from panel A (gray sticks). (C) Overlay of panels A and B. Abbreviation: CRO, chromophore.

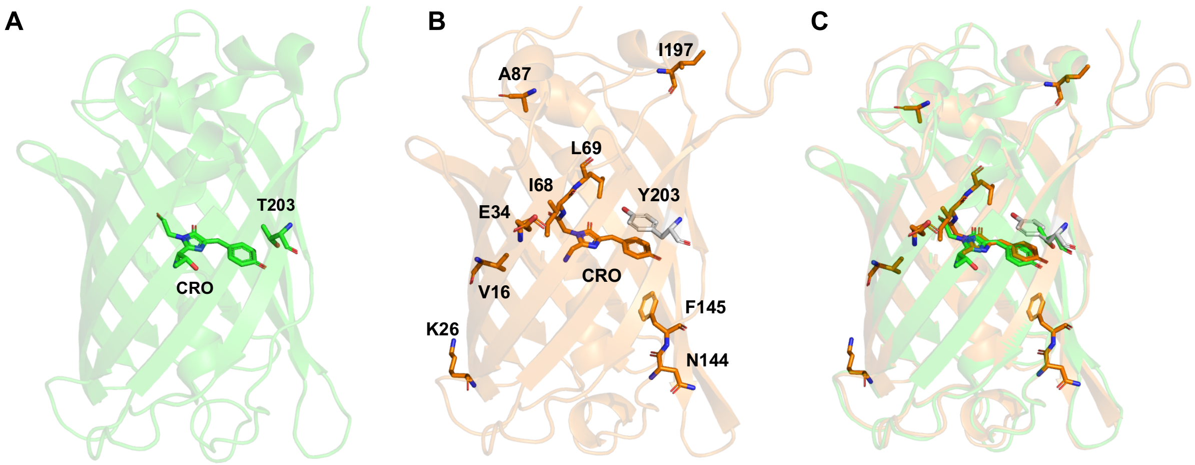


**Figure S35.** (A) Structure of Superfolder GFP (PDB ID: 2B3P) with the chromophore (green sticks) and mutated residues (green sticks) in Table S5 for entry 22. (B) Homology model of OFPxm with the chromophore (orange sticks), all mutated residues (orange sticks), homologous mutated residues from panel A (gray sticks). (C) Overlay of panels A and B. Abbreviation: CRO, chromophore.

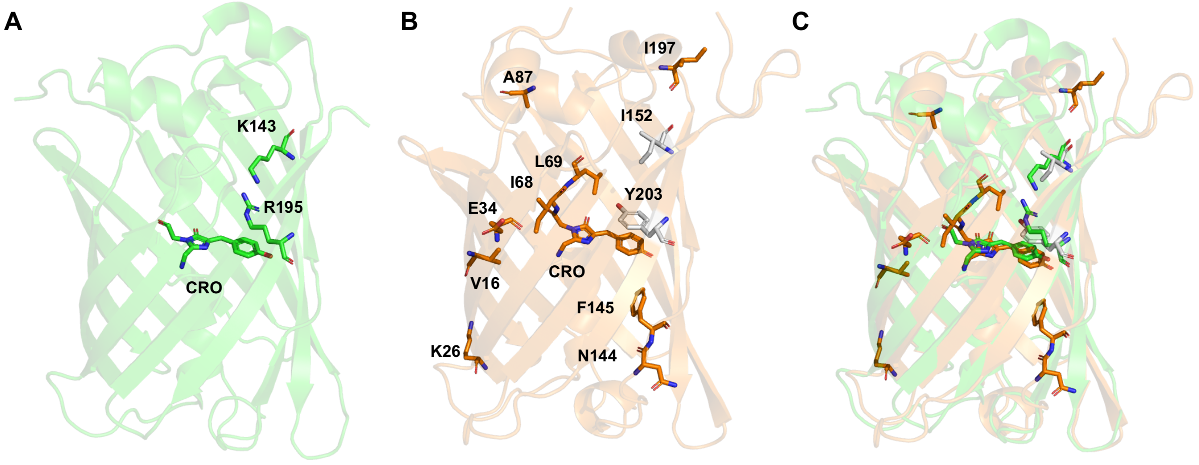


**Figure S36.** (A) Structure of mNeonGreen (PDB ID: 5LTP) with the chromophore (green sticks) and mutated residues (green sticks) in Table S5 for entries 23–26. (B) Homology model of OFPxm with the chromophore (orange sticks), all mutated residues (orange sticks), homologous mutated residues from panel A (gray sticks). (C) Overlay of panels A and B. Abbreviation: CRO, chromophore.

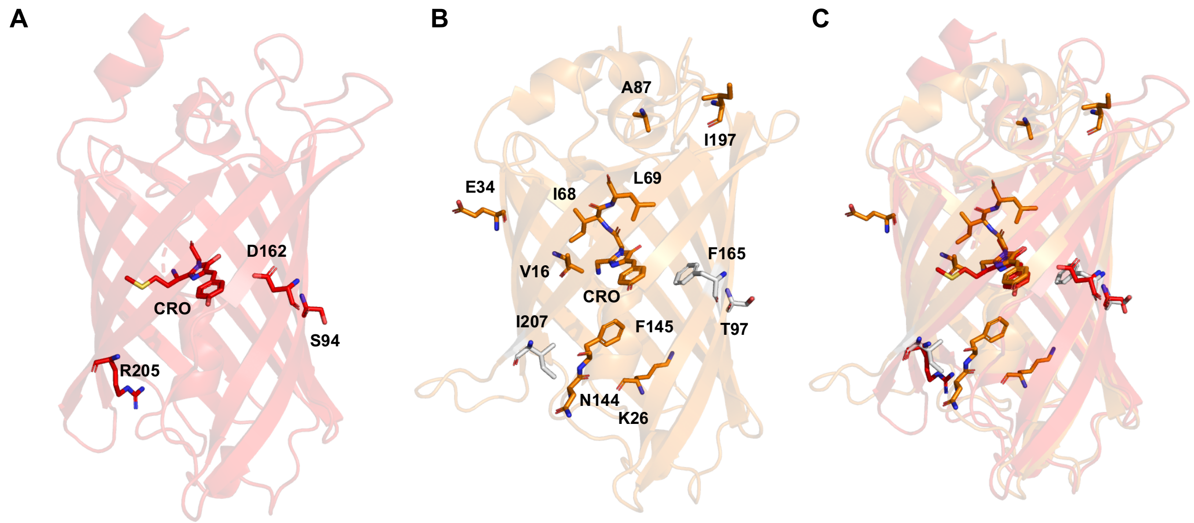


**Figure S37.** (A) Homology model of mBeRFP with the chromophore (red sticks) and mutated residues (red sticks) in Table S5 for entries 27–34. (B) Homology model of OFPxm with the chromophore (orange sticks), all mutated residues (orange sticks), homologous mutated residues from panel A (gray sticks). (C) Overlay of panels A and B. Abbreviation: CRO, chromophore.

### *References*


(S1) McWilliam, H., Li, W., Uludag, M., Squizzato, S., Park, Y. M., Buso, N., Cowley, A. P., and Lopez, R. (2013) Analysis tool web services from the EMBL-EBI. *Nucleic Acids Res. 41*, W597–600.

(S2) Webb, B., and Sali, A. (2016) Comparative protein structure modeling using MODELLER. *Curr. Protoc. Bioinforma. 54,* 5.6.1–5.6.37.

(S3) Wachter, R. M., Yarbrough, D., Kallio, K., and Remington, S. J. (2000) Crystallographic and energetic analysis of binding of selected anions to the yellow variants of green fluorescent protein. *J. Mol. Biol. 301*, 157–171.

(S4) Wang, Q., Byrnes, L. J., Shui, B., Röhrig, U. F., Singh, A., Chudakov, D. M., Lukyanov, S., Zipfel, W. R., Kotlikoff, M. I., and Sondermann, H. (2011) Molecular mechanism of a green-shifted, pH-dependent red fluorescent protein mKate variant. *PLOS ONE 6*, e23513.

(S5) Peng, W., Maydew, C. C., Kam, H., Lynd, J. K., Tutol, J. N., Phelps, S. M., Abeyrathna, S., Meloni, G., and Dodani, S. C. (2022) Discovery of a monomeric green fluorescent protein sensor for chloride by structure-guided bioinformatics. *Chem. Sci. 13*, 12659–12672.

(S6) Heldin, C. H., Johnsson, A., Wennergren, S., Wernstedt, C., Betsholtz, C., and Westermark, B. (1986) A human osteosarcoma cell line secretes a growth factor structurally related to a homodimer of PDGF A-chains. *Nature 319*, 511–514.

(S7) Tutol, J. N., Ong, W. S. Y., Phelps, S. M., Peng, W., Goenawan, H., and Dodani, S. C. (2024) Engineering the ChlorON Series: Turn-on fluorescent protein sensors for imaging labile chloride in living cells. *ACS Cent. Sci. 10*, 77–86.

(S8) Schindelin, J., Arganda-Carreras, I., Frise, E., Kaynig, V., Longair, M., Pietzsch, T., Preibisch, S., Rueden, C., Saalfeld, S., Schmid, B., Tinevez, J.-Y., White, D. J., Hartenstein, V., Eliceiri, K., Tomancak, P., and Cardona, A. (2012) Fiji: an open-source platform for biological-image analysis. *Nat. Methods. 9*, 676–682.
